# Mammary lineage dictates homologous recombination repair and PARP inhibitor vulnerability

**DOI:** 10.1101/2021.05.14.444217

**Authors:** Hyeyeon Kim, Alison E. Casey, Luis Palomero, Kazeera Aliar, Mathepan Mahendralingam, Michael Parsons, Swami Narala, Francesca Mateo, Stefan Hofer, Paul D. Waterhouse, Thomas Kislinger, Miquel A. Pujana, Hal K. Berman, Rama Khokha

## Abstract

It has long been assumed that all normal cells have the same capacity to engage homologous recombination (HR) and non-homologous end joining (NHEJ) to repair DNA double-strand breaks (DSBs), a concept exploited for DNA-damaging chemotherapeutics. We show that mammary epithelial lineage dictates the DSB repair pathway choice. Primary mammary proteomes and DSB repair enumeration by γ-H2AX, Rad51 and DNA-PKc foci reveal that NHEJ operates in all epithelial cells, but high-fidelity HR is restricted to the luminal lineage. This translates to divergent poly (ADP-ribose) polymerase inhibitor (PARPi) vulnerability of mammary epithelial progenitor activity in both mouse and human, irrespective of the *BRCA1/2* status. Proteome-defined lineage-specific signatures correlate to breast cancer subtypes and predict PARPi response of triple-negative human breast cancer xenografts. These intrinsically divergent HR characteristics of mammary cell types underpin a new strategy for identifying PARPi responders.

---

Breast cancer is a heterogenous disease with diverse molecular alterations, histopathological characteristics, response to treatment and patient survival. To understand cellular heterogeneity, early studies of global gene expression profiling of breast tumours have revealed at least five different molecular subtypes (Luminal A, Luminal B, HER2+, Basal-like and Claudin-low)^1,2^, which improved the predictions of prognosis and chemotherapy responses in patients^3^. These subtypes have shown a strong molecular resemblance to distinct epithelial cell types within the normal mammary gland^4^, and hence, each mammary epithelial cell type has been postulated as the precursor cell or the ‘cell-of-origin’ for their corresponding subtype^4^. However, how each cell-of-origin responds to anti-cancer therapies have only begun to be explored^5–7^.

The normal adult mammary epithelium is a bilayered ductal structure composed of two major lineages, luminal and basal, each of which contains a spectrum of populations of varied differentiation potential and cellular states^8,9^. Fluorescence-activated cell sorting (FACS) can isolate three phenotypically distinct mammary epithelial cell populations: basal, luminal progenitor, and luminal mature^10–12^ which are essentially the same populations pinpointed as the major mammary epithelial cell types from unbiased, marker-free classifications from the recent single-cell RNA sequencing studies^9,13,14^. In general, the luminal progenitor population is enriched for luminal lineage-restricted progenitor cells that sustain the luminal layer^15,16^, and is largely hormone receptor negative while the luminal mature population is enriched for more differentiated, hormone receptor positive cells that lack clonogenic capacity^17^. Unlike the two luminal populations, the FACS-defined basal population is highly plastic and heterogenous^8,18^, harbouring mixed cell populations of differentiated myoepithelial cells, basal-restricted progenitors, and possibly, rare bipotent stem cells^9,19^. These populations have many biological distinctions which can be elucidated to improve their targeting as cell-of-origin in breast cancer.

In recent years, these three main epithelial cell types have been subjected to bulk transcriptome^8,14,20,21^, epigenome^20,22^, and proteome^5,6^ profiling in order to identify unique biological features that distinguish one cell type from the others, and possibly exploit targeting of the putative cell-of-origin for that breast cancer subtype. Our recently published proteomic datasets on mouse^5^ and human^6^ mammary epithelial cells, led us to identify differential DDR protein expression across the three main epithelial lineages that we posit leads to intrinsically divergent cellular activity. We show that the functional capacity of HR repair differs across the mammary lineages and results in a differential response to PARPi, a class of cancer drugs that target HR deficiency. To our knowledge, this study is the first to identify cell lineage as a non-mutational determinant of PARPi sensitivity, and our proteome-based mammary lineage-specific signatures may clinically inform the most appropriate intervention strategy to enhance PARPi efficacy.

## Results

### Normal mammary epithelial populations have disparate DNA damage response

We interrogated our published normal mouse mammary proteome resource^5^ to identify biological functions uniquely upregulated in basal (<u>Lin^−^CD24^+^CD49f^hi^</u>), luminal progenitor (<u>Lin^−^CD24^+^CD49f^lo^Sca1^−^CD49b^+^</u>), and luminal mature (<u>Lin^−^CD24^+^CD49f^lo^Sca1^+^CD49b^+/−^</u>) populations^10,12^, all FACS-purified from virgin female wild-type mice (**Fig. 1a**). Gene set enrichment analysis^23^ revealed 57, 63, and 8 pathways to be uniquely upregulated (false discover rate (FDR) <0.01) in basal, luminal progenitor, and luminal mature populations, respectively (**Fig. 1b** and **Supplementary Table 1**). Notably, one-third (23/63) of all enriched pathways in the luminal progenitor were related to the DNA damage repair (DDR) function (**Fig. 1c**). Most known breast cancer susceptibility genes participate in DDR^24^, such as *ATM*, *CHEK2, MRE11*, *BRCA1*, *BRCA2*, *RAD51* paralogues, where a functional deficiency in HR repair (e.g. deleterious mutations in *BRCA1*/*2*) is prevalent in up to 22% of all breast cancer cases^25^. Of the 156 proteins known to participate in the DSB repair pathway (R-HSA-5693532; Reactome Database ID Release 63), 32 were detected in our mouse mammary proteomic dataset and about two-thirds of them were more abundant in the luminal progenitor population (**Fig. 1d**). We therefore decided to investigate the physiological DDR across mammary epithelial lineages.

**Fig. 1.**
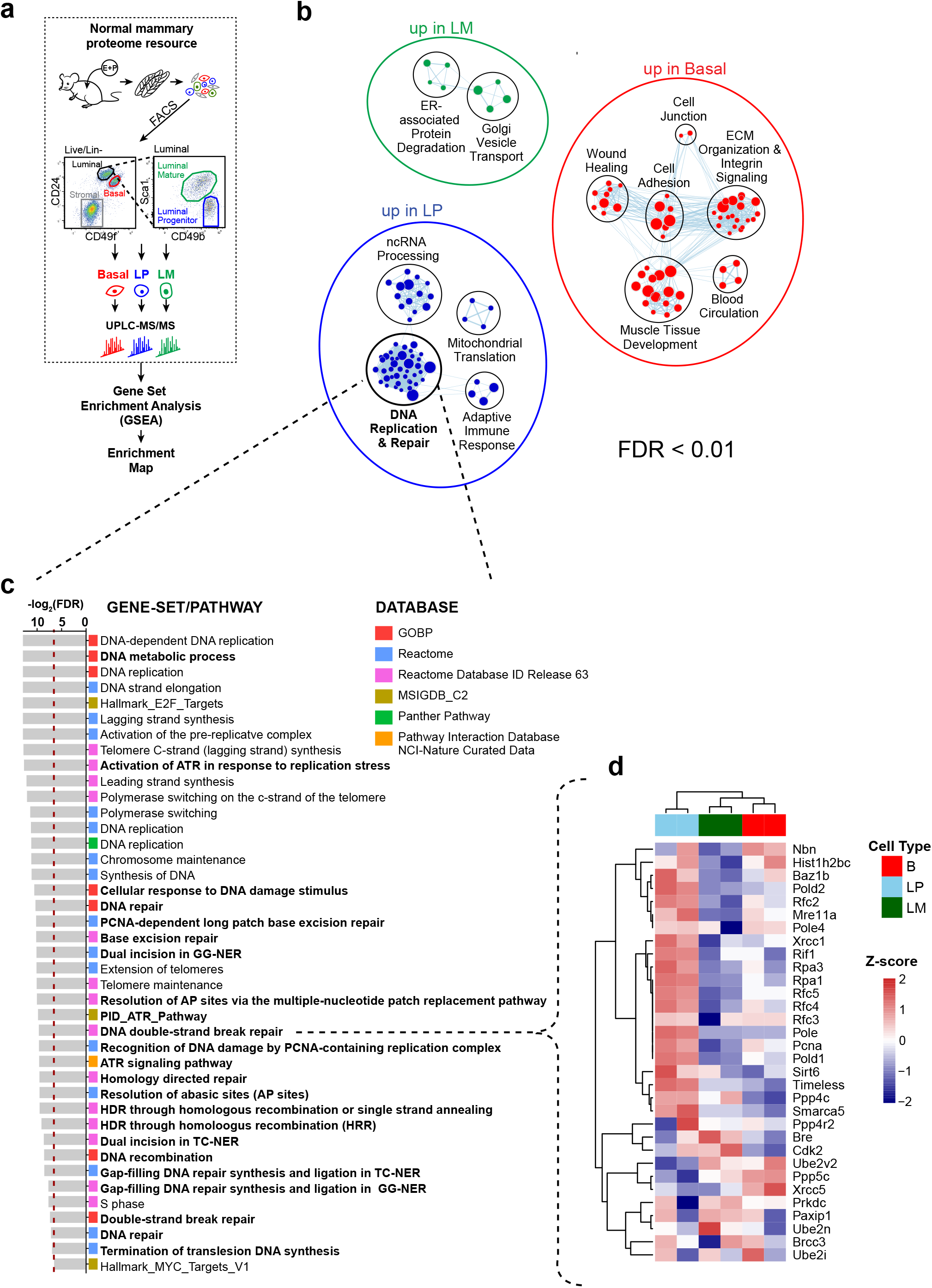
Global proteomics of normal mouse mammary epithelial populations reveal their specific biological programs. **(a)** A schematic diagram illustrating interrogation of our previously published normal mouse mammary proteomic dataset. Global proteomic profiles of FACS-purified basal (red; Lin^−^ CD24^+^CD49^hi^), luminal progenitor (LP; blue; Lin^−^CD24^+^CD49^lo^Sca1^−^CD49^+^), or luminal mature (LM; green; Lin^−^CD24^+^CD49^lo^Sca1^+^CD49^+/−^) epithelial population cells from the mammary glands of wild-type female mice under sex-hormone stimulation (estrogen plus progesterone; E+P) were subjected to pathway enrichment analysis using Gene Set Enrichment Analysis (GSEA). The resulting enriched pathways (FDR<0.01) for each population were visualized into a network by using Enrichment Map from Cytoscape. **(b)** Enrichment map illustrating clusters of gene-sets/pathways (nodes) significantly upregulated in the basal, luminal progenitor (LP), or luminal mature (LM) population compared to the other two as determined by GSEA (FDR<0.01). Each cluster was manually annotated with a common biological theme. **(c)** Each pathway is colour-coded by corresponding curated database, and pathways related to DNA damage repair are bolded. The red-dashed line indicates the −log_2_(FDR) cut-off. **(d)** Heatmap showing unsupervised hierarchical clustering of DDR protein abundance in each mammary epithelial population (n=2; pooled from multiple mice) curated in the “DNA double-strand break repair” pathway (R-HSA-5693532; Reactome Database ID Release 63).

Neutral comet assay measures Olive tail moment as a cellular feature of DSBs in individual cells. FACS-purified mammary populations from un-irradiated wild-type mice revealed a longer Olive tail moment in the luminal progenitor compared to the luminal mature or basal cells indicating the higher homeostatic level of DSBs in the luminal progenitor population (**Fig. 2a**). Histone H2AX phosphorylation, γ-H2AX, marking of DSBs is an early event in DDR which acts as a scaffold for the rapid recruitment of repair proteins to the site of DSBs for efficient DNA repair^26^. We optimized the irradiation dose (2 to 6 Gy) and time-points (0.5 to 24 hours) to map *in situ* DNA damage response activation in the mammary gland through staining for γ-H2AX and 53BP1 foci formation (**Extended Data Fig. 1**). We then quantified the kinetics of γ-H2AX in the three populations by intracellular flow cytometry (**Fig. 2b** and **Extended Data Fig. 2a-e**) of the dissociated mammary gland. Irradiation-induced γ-H2AX peaked at 0.5 h and resolved within 24-48 h in all three mammary epithelial populations, yet individual populations displayed specific kinetics of γ-H2AX foci formation and resolution over the time course (**Fig. 2b**). The three epithelial populations had similar proportions of γ-H2AX^+^ cells at the early time-points (**Extended Data Fig. 2b**), however, luminal progenitors had the highest absolute number of γ-H2AX^+^ cells (**Fig. 2c**). Furthermore, the median fluorescence intensity of γ-H2AX was highest in these cells at the early time-points including the baseline un-irradiated state (**Fig. 2c**). Likewise, punctate γ-H2AX foci visualized by tissue immunofluorescence were primarily detected in proliferating, progesterone receptor negative luminal (Ki67^+^PR^−^K14^−^) cells which characterize luminal progenitors^12,27^ **(Fig. 2d** and **Extended Data Fig. 2f)**. Therefore, luminal progenitor population distinguished itself by markedly higher DNA damage response.

**Fig. 2.**
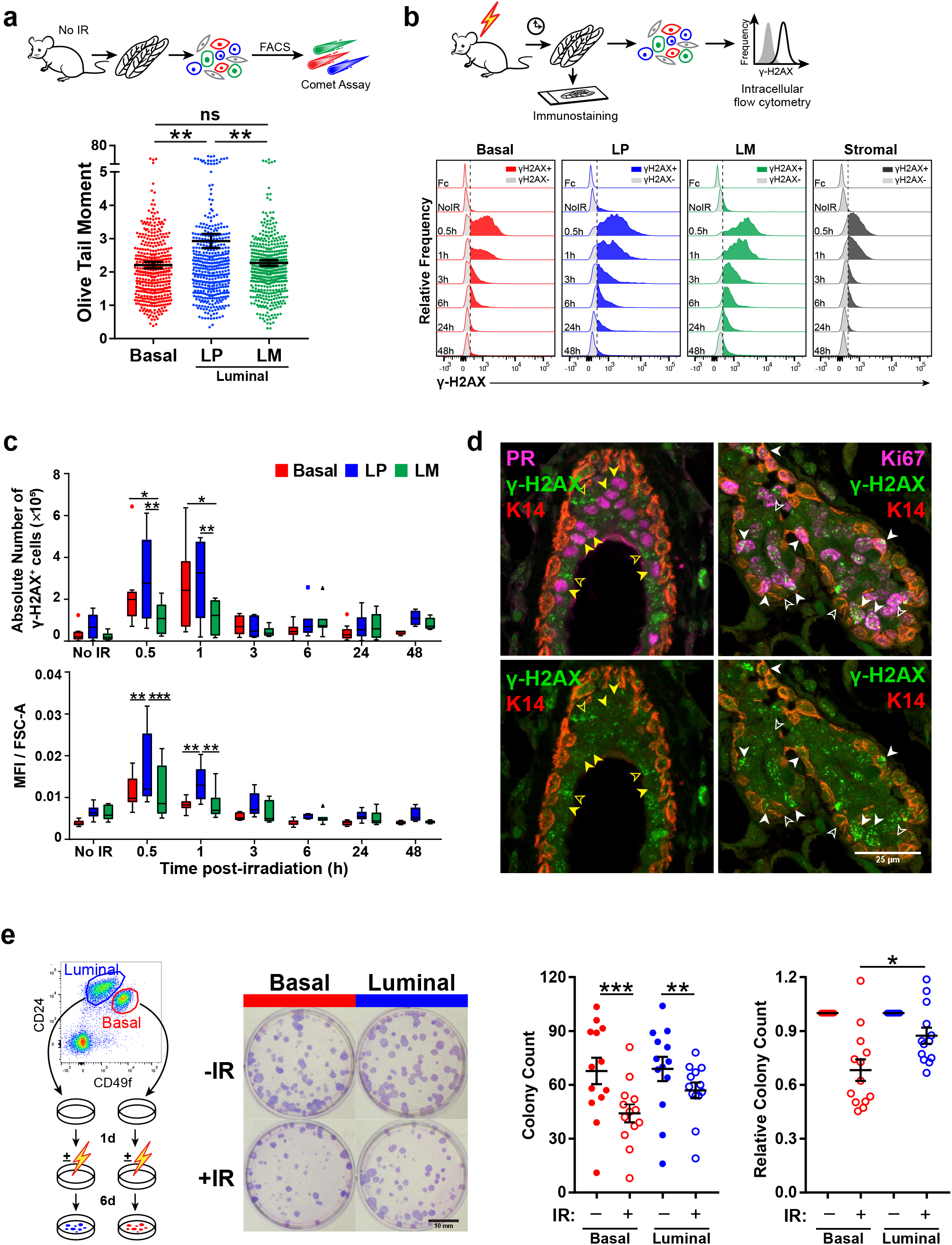
Differential DNA damage response capacity across mammary epithelial populations after irradiation. **(a)** Workflow of neutral comet assay. Scatter plot showing Olive tail moment on sorted cells from basal (red), luminal progenitor (blue; LP), and luminal mature (green; LM) populations from un-irradiated, wild-type female mice. A total of ∼450 comets were scored per population (100-130 comets/population/mouse; n=4 mice). Analysis by one-way ANOVA with Tukey’s multiple comparison test was used. **P<0.01. ns, not significant. **(b)** Workflow depicting intracellular flow cytometry of γ-H2AX on single cells from the mammary gland, and of immunofluorescence staining on the mammary tissue sections following 6 Gy *in vivo* irradiation. Representative flow cytometry histograms of γ-H2AX recruitment in basal, LP, LM, and stromal populations at indicated time-points post-irradiation. The dotted line delineates γ-H2AX positivity (γ-H2AX^+^ cells) from the background (γ-H2AX^−^ cells) based on the Fc control. **(c)** Boxplots showing median fluorescence intensity (MFI) in γ-H2AX^+^ cells normalized by cell size (FSC-A; see Extended Data Fig. 2d,e) and absolute number of γ-H2AX^+^ cells in each population (n=4-10 mice per time-point). Two-way ANOVA with Tukey’s multiple comparison test was applied. *P<0.05, **P<0.01, ***P<0.001. **(d)** Z-projected confocal immunofluorescence images of γ-H2AX (green), basal cytokeratin (K14; red), and progesterone receptor or proliferation (PR or Ki67; magenta) in mammary tissue at 3 h post-irradiation. Filled yellow arrowheads indicate PR^+^ cells and hollow indicate PR^−^ cells. Filled white arrowheads indicate Ki67^+^ cells and hollow indicate Ki67^−^ cells. **(e)** Workflow of colony forming cell (CFC) assay on FACS-purified basal (red) and total luminal (blue) cells after *in vitro* irradiation (3 Gy). Basal and luminal progenitors give arise to corresponding colonies. Luminal mature cells lack clonogenicity in this media condition. Representative images of basal and luminal colonies with/without irradiation (IR) after 7 days of culture. Scatter plots showing the absolute counts of basal (red) and luminal (blue) colonies with/without IR (left; n=13; paired *t*-test). Relative colony count of irradiated samples to the paired un-irradiated sample are also shown on right (n=13; unpaired *t*-test). All data represent mean ± SEM. *P< 0.05, **P<0.01, ***P<0.001.

To assess the functional outcome of a differential DNA damage response, FACS-purified basal and total luminal (<u>Lin^−^CD24^+^CD49f^lo^</u>) lineages were cultured in a colony forming cell (CFC) assay to measure lineage-restricted progenitor activity and clonogenic survival with or without *in vitro* irradiation (**Fig. 2e**). The progenitor capacity of both lineages declined after irradiation as anticipated, yet significantly more luminal colonies survived than the basal colonies (P<0.05; **Fig. 2e**), indicating that luminal progenitors are more radioresistant than basal progenitors. Collectively, these data show that luminal progenitors are physiologically more susceptible to DSBs, however, their ability to elicit potent activation of DNA damage response leads to better survival compared to the basal progenitors.

### Proliferating basal cells do not engage HR upon genotoxic insult

DSBs are the most lethal forms of DNA damage, and thus accurate DSB repair is critical to prevent genetic alterations and genomic instability. HR repair uses the undamaged sister chromatid as a template to accurately repair the DSB region, whereas NHEJ directly ligates the two DSB ends without any guidance from a homologous DNA sequence and is thus more error-prone. We assessed HR and NHEJ activities at the single-cell level using Amnis imaging flow cytometry, which has dual features of flow cytometry and fluorescence microscopy that enable capturing of spatial information of fluorescence signals. High-throughput quantification of nuclear repair foci was performed in each mammary population as defined by their established cell surface markers (**Extended Data Fig. 3a**). Nuclear Rad51 foci and DNA-PKcs (phospho-S2056; pDNA-PKcs) provided functional readouts for HR and NHEJ, respectively (**Fig. 3a** and **Extended Data Fig. 3b**). Before irradiation, the highest proportion of Rad51^+^ cells was in luminal progenitors (18%) followed by the luminal mature population (10%), while Rad51 foci were rarely found in basal cells (1.2%; **Fig. 3b**). This equated to up to 15-fold higher HR activity in luminal progenitor versus the basal lineage. At 3h post-irradiation, the proportion of Rad51^+^ cells increased in all populations as anticipated, and the luminal lineage still showed a ∼4.5-fold higher Rad51^+^ proportion compared to basal (P<0.0001; **Fig. 3b**). Further, luminal progenitors harboured the highest absolute number of Rad51^+^ cells at baseline and post-irradiation (**Extended Data Fig. 3c**). Conversely, the largest proportion of pDNA-PKcs^+^ cells were found in luminal mature (34% at baseline and 54% at 3h post-irradiation) followed by luminal progenitor (20% and 33%) and basal (5% and 24%) cells, although basal cells displayed the highest fold increase in NHEJ activity upon irradiation (**Fig. 3c** and **Extended Data Fig. 3d**). Collectively, these data demonstrate that NHEJ is universally utilized by all mammary epithelial cells, but HR is predominant in the luminal lineage, especially in the luminal progenitor population.

**Fig. 3.**
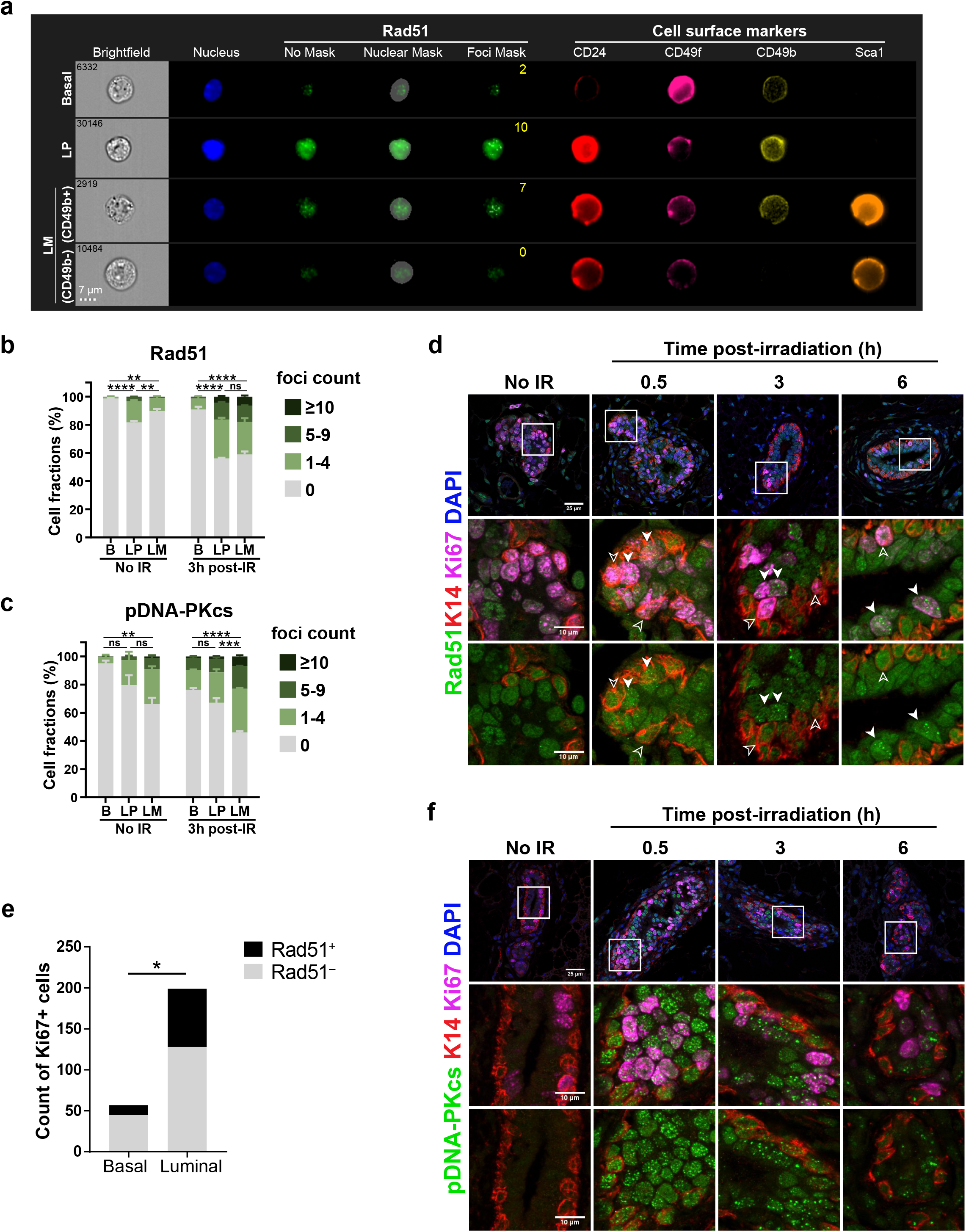
Homologous recombination (HR) repair is intrinsically different between luminal progenitors and basal cells. **(a)** A panel of representative ImageStream®^X^ images displaying Rad51 foci in the basal, luminal progenitor (LP), and luminal mature (LM) populations as marked by a set of established cell surface markers. Foci counts, as determined by IDEAS^®^ software, are indicated in yellow in the Rad51 ‘Foci Mask’ column. **(b,c)** Bar graphs summarizing proportion of cells displaying 0, 1-4, 5-9, or ≥ 10 foci of (b) Rad51 or (c) DNA-PKcs (phosho-S2056; pDNA-PKcs) in each population before or 3 h-post irradiation. (n=3-4 mice per treatment group). Data represent mean ± SEM and p-values determined by one-way ANOVA with Tukey’s multiple comparison test Rad51^+^ or pDNA-PKcs^+^ cells (i.e. cells displaying ≥ 1 focus) across all 3 mammary populations. **P<0.01, ***P<0.001, ****P<0.0001. ns, not significant. **(d,f)** Z-projected confocal immunofluorescence images of (d) Rad51 and (f) pDNA-PKcs co-stained with basal cytokeratin (K14) and proliferation (Ki67) markers on paraffin-embedded mammary tissues harvested from virgin female mice with/without *in vivo* irradiation. Filled white arrowheads indicate proliferating Rad51^+^ cells, and hollow arrowheads indicate proliferating Rad51^−^ cells. **(a) (e)** Absolute counts and proportion of proliferating/Ki67^+^ cells exhibiting Rad51^+^ (≥2 foci) in basal (K14^+^) or luminal (K14^−^) cells at 3 h post-irradiation by immunofluorescence staining. The number of Rad51 foci per nucleus was determined by ImageJ. P-value was calculated by Fisher’s exact test. *P<0.05.

We then delineated cell proliferation status in relation to repair since DSB repair is tightly regulated by the cell-cycle; HR is restricted to cells in late S and G2 phases whereas NHEJ is predominantly used during the G1 phase of the cell cycle^28^. Our analysis of HR kinetics, as determined by *in situ* immunofluorescence staining of the mammary gland, showed that Rad51 foci formed strictly in proliferating (Ki67^+^) cells as early as 0.5 h, and progressed to larger and punctate foci by 3-6 h post-irradiation (**Extended Data Fig. 4a**, filled arrowheads). Rad51 foci were present in proliferating luminal (Ki67^+^K14^−^), but largely absent in proliferating basal cells (Ki67^+^K14^+^; P<0.05; **Fig. 3d,e** and **Extended Data Fig. 4a**). Further luminal characterization by co-staining of the PR showed that Rad51 foci were predominantly found in luminal progenitors (K14^−^PR^−^Ki67^+^) rather than luminal mature cells (K14^−^PR^+^Ki67^−^; **Extended Data Fig. 4b,c**). Conversely, pDNA-PKcs^+^ foci were seen in both compartments at all time-points and mostly in Ki67^−^ cells, an observation consistent with the known cell-cycle regulation (**Fig. 3f**). Hence, among the proliferating mammary epithelial cells, only those with luminal progenitor characteristics displayed the HR hallmark. In addition to, and perhaps prior to, cell cycle regulation, DSB repair pathway choice is directed by cell lineage in the normal adult mammary gland.

### Basal cells are more sensitive to PARPi than luminal progenitors irrespective of BRCA status

PARPi have proven of immense clinical value in treating cancers with HR defects including those arising in *BRCA*-mutation carriers^29^. Although PARPi were developed to inhibit the catalytic activation of PARP, studies showed most potent PARPi exert their therapeutic effect by “trapping” PARP1/2 at single-strand DNA breaks^30,31^, creating a PARP-DNA complex that induces replication fork collapse and *de novo* DSBs^32^. As HR repair is essential to resolve these PARPi-induced DSBs^30^, cells defective in the HR pathway, such as those harbouring deleterious *BRCA* mutations, are hypersensitive to PARPi due to a synthetic lethal interaction^33,34^. Since we observed differential HR engagement in the two normal mammary lineages, we assessed the effect of PARPi on progenitor activity using CFC assays as a readout (**Fig. 4a**). Luminal and basal progenitors exhibited strikingly divergent vulnerability to PARPi. Basal CFCs decreased precipitously in a dose-dependent manner in response to olaparib and talazoparib, whereas luminal CFCs were comparatively resistant to these drugs (**Fig. 4b**). Meanwhile, treatment with KU-57788, a DNA-PK inhibitor that targets NHEJ, showed no apparent lineage bias (**Fig. 4b**).

**Fig. 4.**
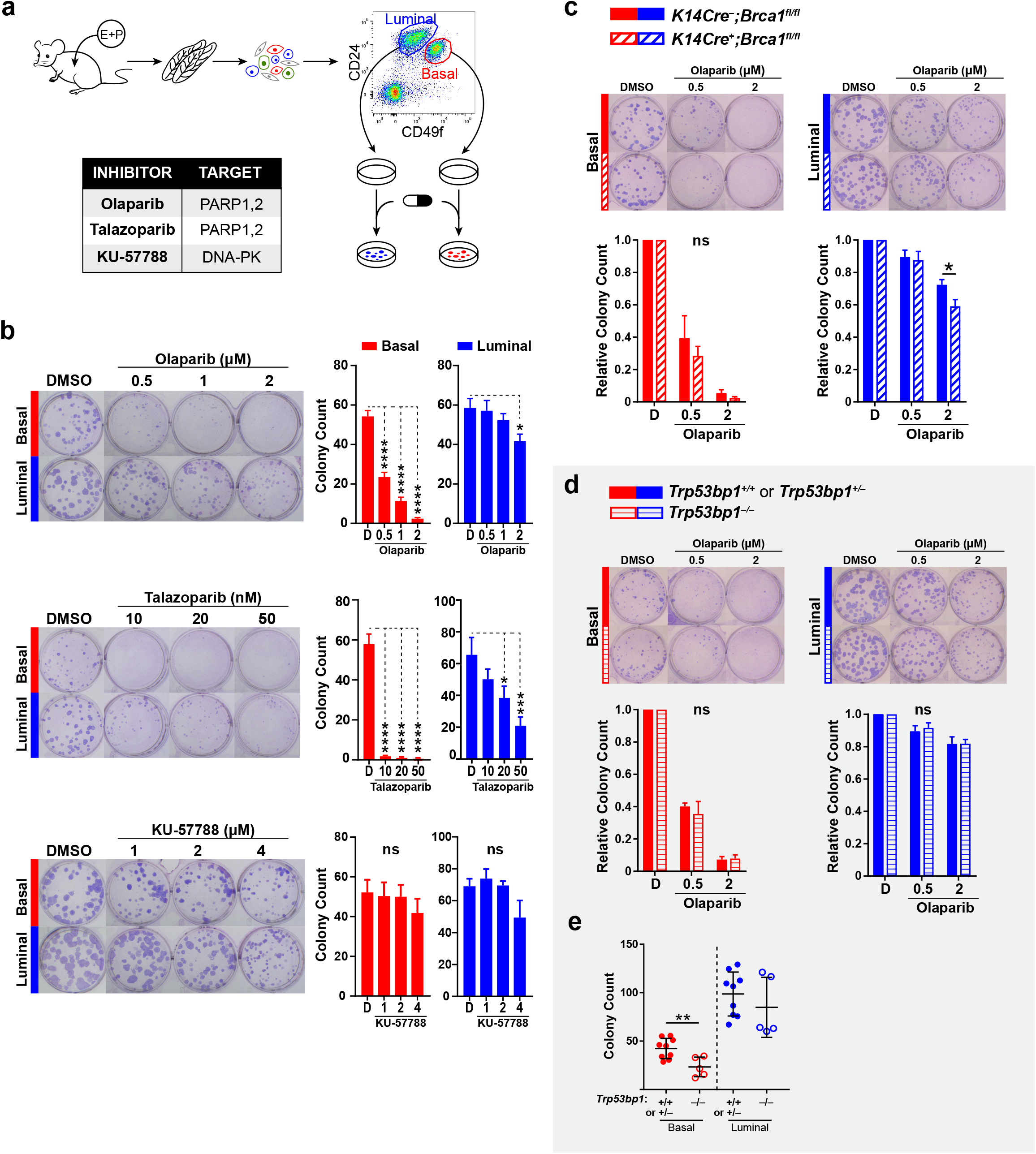
Luminal and basal progenitors exhibit divergent PARPi vulnerability. **(a)** Workflow of colony forming cell (CFC) assay from FACS-purified basal (<u>Lin^−^ CD24^+^CD49f^hi^</u>; red) and total luminal (<u>Lin^−^CD24^+^CD49f^lo^</u>; blue) fractions to evaluate basal and luminal progenitor capacities in response to the DDR-targeting drugs listed in the table. **(b)** Representative images of basal and luminal colonies treated with olaparib, talazoparib, KU-57788, or DMSO alone (vehicle control). Bar graphs show the CFC count for basal (red) and luminal (blue) populations at the indicated drug concentrations after 7 days of culture (n=5-8 mice). One-way ANOVA with Dunnett’s multiple comparisons test was performed. *P<0.05, ***P<0.001, ****P<0.0001. **(c,d)** Olaparib sensitivity of the mammary lineages from (c) *Brca1*-deficient mice (*K14-cre^+^;Brca1^fl/fl^*; n=6) and littermate controls (n=5-7) or from (d) *Trp53bp1*-deficient (*Trp53bp1*^−/–^; n=5) and littermate controls (n=9) using the above CFC workflow. Bar graphs show the number of basal or luminal colonies relative to DMSO control after olaparib treatment. Data represent mean ± SEM. Two-way ANOVA with Sidak’s multiple comparison’s test was performed. *P<0.05. ns, not significant. **(a) (e)** Scatter plot demonstrating the absolute colony count of un-irradiated, DMSO-treated basal and luminal colonies from littermate control (*Trp53bp1*^+/+^ or *Trp53bp1*^+/−^; n=9) or experimental (*Trp53bp1*^−/–^; n=5) cohorts. All data represented as mean ± SEM. P-value was determined by t-test. **P<0.01.

We investigated whether the loss of *Brca1* would sensitize luminal progenitors to olaparib since a synthetic lethal interaction occurs between the PARPi and *BRCA* mutations. *K14-cre*^+^*;Brca1^fl/fl^* mice lack *Brca1* in the entire mammary epithelium due to K14 (cytokeratin 14) promoter activity during embryonic mammary development^35^. A *Brca1* deficiency was expected to increase olaparib sensitivity, however, *K14-cre^+^;Brca1^fl/fl^* luminal CFCs were only moderately more sensitive compared to their wild-type littermates (*K14-cre*^−^*;Brca1*^F/F^; **Fig. 4c**). The loss of *Brca1* had no effect on basal CFCs, which remained hypersensitive to olaparib (**Fig. 4c**). We also employed transformation-related p53-binding protein 1-null (*Trp53bp1*^−/–^) mice in which cells can no longer repair DSBs via NHEJ^28^. *Trp53bp1*-deficient basal and luminal CFCs showed no greater olaparib sensitivity compared to their littermate controls, but still retained the same lineage-specific PARPi vulnerability (**Fig. 4d**). We also noted lower baseline clonogenic potential in the untreated *Trp53bp1*-deficient basal CFCs compared to *Trp53bp1*^+/+^ or *Trp53bp1*^+/−^ controls (**Fig. 4e**), suggesting that NHEJ-defective basal progenitors are more sensitive to replication stress in the growth factor enriched CFC assay conditions. Collectively, by exploiting PARPi-mediated synthetic lethality, we confirmed functional differences in HR capacity between the two normal mammary lineages in which basal progenitors are intrinsically sensitive, but luminal progenitors are largely resistant to PARPi, even with a homozygous deletion of *Brca1*.

We examined whether this lineage-dependent vulnerability to PARPi is conserved in humans. Unlike mouse, primary normal human breast progenitor cells give rise to 3 morphologically distinct colonies *in vitro*: myoepithelial-restricted, luminal-restricted, and bipotent, which are indicative of the differentiation capacity of their respective progenitor cell populations^36^. Myoepithelial colonies represent more differentiated myoepithelial progenitors in the basal (<u>Lin^−^ EpCAM^lo^CD49f^hi^</u>) population, luminal colonies arise from the luminal progenitors (<u>Lin^−^ EpCAM^hi^CD49f^hi^</u>), and bipotent colonies are mostly enriched in the basal fraction^37^. On the other hand, the luminal mature (<u>Lin^−^ EpCAM^hi^CD49f^lo^</u>) population lacks clonogenicity in conventional CFC culture conditions^37,38^ (**Fig. 5a**). Primary breast tissues from 20 patients who underwent prophylactic mastectomy, including *BRCA* wild-type, *BRCA1*- and *BRCA2*-mutation carriers (n=5, 9, 6 respectively; 30-64 years old), were dissociated into single cells for phenotypic flow cytometry and matched CFC scoring (**Fig. 5a** and **Extended Data Fig. 5a-c**). We observed cellular heterogeneity across all patients as well as within *BRCA* cohorts (**Extended Data Fig. 5a)**. The control (DMSO) CFC score for each colony type generally reflected the proportions of the epithelial cells in their corresponding flow profiles (**Extended Data Fig. 5b**). We found that olaparib predominantly affected basal progenitor capacity; abrogating 13% of myoepithelial (P<0.05, ratio paired t-test) and 47% of bipotent colonies (P<0.0001). In contrast, luminal colonies remained largely resistant in all patients (**Fig. 5b**). Although higher olaparib sensitivity was observed in all three colony types generated from *BRCA* carrier samples (P<0.05; Two-way ANOVA; **Fig. 5c**), the colony type remained the major factor in olaparib response, with luminal lineage being more resistant than basal, even when *BRCA* mutations are present (P<0.0001; Two-way ANOVA; **Fig. 5c**). Thus, mammary lineage is the primary determinant for olaparib vulnerability in both the mouse and human mammary epithelium.

**Fig. 5.**
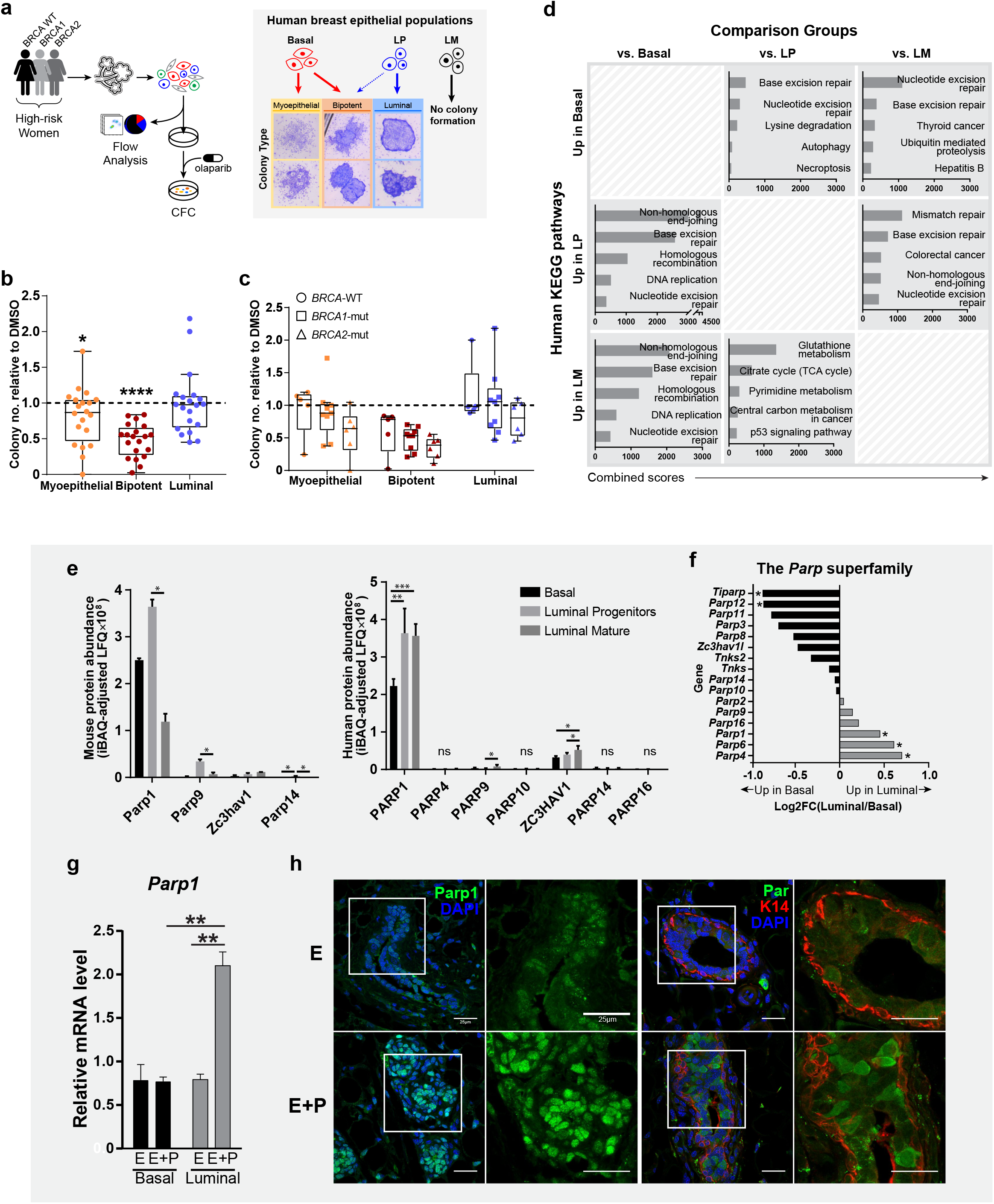
Olaparib reduces human mammary basal progenitor capacity. **(a)** Workflow of the 21 primary human breast specimens from *BRCA* wild-type, *BRCA1* or *BRCA2*-mutated carriers (n=5, 9, 6, respectively) for flow cytometry and olaparib testing via colony forming cell (CFC) assay (left). Schematic diagram illustrating human mammary epithelial populations and representative images of three types of colony that arise from corresponding progenitor activities (right). **(b)** Boxplot summarizing growth of myoepithelial, bipotent or luminal colonies compared to DMSO vehicle control (horizontal dotted line), in all primary breast specimens (n=20). P-value for colony growth in olaparib vs DMSO was calculated by ratio paired t-test. *P<0.05, ****P < 0.0001. **(c)** Boxplot summarizing the effect of olaparib on the growth of each colony type, according to BRCA mutation status (*BRCA* wild-type, n=5; *BRCA1*-mut, n=9; *BRCA2*-mut, n=6), compared to DMSO control (horizontal dotted line). Whether the two independent factors, colony type and BRCA mutation status, affected differential colony growth was tested by performing two-way ANOVA. The colony type significantly affected colony growth upon olaparib treatment (***p<0.001). The BRCA status significantly affected colony growth upon olaparib treatment (*p<0.05). No interaction was found between these two factors (p≤0.9754). Post-hoc analysis by Tukey’s multiple comparisons test showed no differences in colony growth of each colony type based on BRCA mutation status (p-values ranging from 0.24-0.98). **(d)** Significantly upregulated (q<0.05) human DDR proteins found in three different pairwise comparisons between in basal, luminal progenitor (LP), and luminal mature (LM) populations (1 Data Fig. 6b) were subjected to pathway analyses by Enrichr. The top 5 human KEGG pathways for every pair were determined based on the ‘combined’ score from Enrichr. **(e)** Protein abundance values of detected Parp/PARP members from mouse (left; n=2, each pooled from multiple mice) or human (right; n=6) mammary proteome^5^. Differential abundance of each PARP member across different mammary populations was determined by using one-way ANOVA with Tukey’s multiple comparisons test. *P<0.05, **P<0.01, ***P<0.001. ns, not significant. **(f)** Mean log2 fold-change (log2FC) of mRNA expression level for 16 (out of 17) members of the Parp superfamily (Parp15 not detected) in the luminal versus basal lineage determined from a published microarray analysis^42^ on FACS-purified luminal and basal cells from female virgin mice under ovarian hormone-stimulated (estrogen plus progesterone; E+P) state (n=4/group). Asterisk indicates q-value < 0.05. **(g)** *Parp1* mRNA levels measured by qRT-PCR in FACS-purified primary mouse basal and total luminal cells from mice treated with defined sex hormones (estrogen; E or estrogen plus progesterone; E+P). Expression is normalized to β-actin (endogenous control) and relative to E basal. P-values were determined by using unpaired *t*-test. **P < 0.01. **(h)** Co-immunostaining of cytokeratin 14 (K14) and PARP1 or PAR on mouse mammary tissue sections from virgin female mice under defined sex-hormone treatment: estrogen (E) or estrogen plus progesterone (E+P). Scale bar = 25 μm for all panels.

### Luminal lineage exhibits higher baseline and progesterone-induced Parp1 activity

To understand the human DDR proteomic landscape, we data-mined our recently published global human mammary proteome^6^ that has profiled the three mammary populations (basal, luminal progenitors, luminal mature) from 6 normal, premenopausal breast tissues. 127 of the 276 human DDR genes curated by Knijnenburg *et al*^39^ were detected in the human proteome (**Extended Data Fig. 6a)**. Unsupervised hierarchical clustering revealed that these human DDR proteins clustered based on the mammary cell type, and strikingly, most of them appeared to be less abundant in the basal population (**Extended Data Fig. 6a**). Pairwise comparisons uncovered differentially expressed DDR proteins between the cell types, and significantly upregulated DDR proteins from each comparison (q<0.05) were subjected to pathway analysis using Enrichr^40,41^ (**Extended Data Fig. 6b** and **Supplementary Table 2**). KEGG pathways relating to NHEJ and HR were enriched in human luminal progenitor and luminal mature populations, whereas the basal population was enriched for single-strand break repair pathways, such as nucleotide excision repair and base excision repair (**Fig. 5d**). This illustrates the heterogenous DDR proteomic landscape of human breast epithelium as observed in the mouse.

We sought the molecular mechanism underlying divergent PARPi response of mammary clonogenicity by examining PARP superfamily members in our global mammary datasets including mouse transcriptomes, mouse proteomes and human proteomes. PARP1 is the major target of PARPi, and its genetic deletion confers high PARPi resistance^30^. Of all detected PARPs in the proteomes, PARP1 was the most abundant in both mouse and human datasets (**Fig. 5e)**. We also noted that human PARP1 was ∼1.6-fold higher in luminal progenitor and luminal mature populations compared to basal (**Extended Data Fig. 6c)**. In mouse microarray data^42^, 16 of 17 *Parp* genes were found, including *Parp1*, *Parp4*, and *Parp6*, which were significantly higher in total luminal, while *Tiparp* (*Parp7*) and *Parp12* were higher in the basal cells (q<0.05; **Fig. 5f**). Progesterone is known to expand the number of adult stem/progenitor cells in the mammary gland^43,44^, and induced a 2.5-fold increase in *Parp1* mRNA level only in the luminal cells (P<0.01; **Fig. 5g**). Co-immunostaining of Par and K14 on the normal, un-irradiated mouse mammary gland following progesterone treatment, provided *in situ* confirmation of increased Parp1 activity in the luminal versus basal layer of the mammary gland (**Fig. 5h**). Despite having more targets for PARPi in luminal cells, their clonogenic capacity remains intact likely due to their ability to carry out the HR repair. The basis for lineage differences in PARP proteins and activity requires further study.

### Proteome-defined lineage IDs inform breast cancer subtypes and PARPi response

We next asked whether lineage identification (ID) can serve as a non-mutational predictor of olaparib sensitivity in breast cancers. We first constructed a set of human proteome signatures (>3-fold enrichment; **Supplementary Table 3**) as IDs for the basal, luminal progenitor and luminal mature populations, and then examined their association with breast cancer subtypes from the METABRIC dataset^45^. The human basal proteomic ID showed the highest correlation to the mesenchymal ‘Claudin-low’ subtype; the luminal progenitor ID was most closely linked to ‘Basal-like’, a subtype that arises frequently in *BRCA1* breast cancers^46,47^; and the luminal mature ID was highly correlated to the most commonly diagnosed, hormone-receptor-expressing ‘Luminal A’ subtype (**Fig. 6a** and **Extended Data Fig. 7a,b**), extending the previously proposed cell-of-origin/subtype model based on transcriptome data^4^ to hold true at the protein level. The same association was seen with mouse mammary proteome-generated lineage IDs (**Extended Data Fig. 8a,b** and **Supplementary Table 4**). Additionally, both human and mouse luminal progenitor IDs positively correlated with somatic mutation profile #3^48^, which is associated with HR deficiency and deleterious *BRCA1/2* mutations, whereas the luminal mature IDs showed an inverse correlation and the basal IDs showed no association with defective HR (**Fig. 6b**, **Extended Data Fig. 8c** and **Supplementary Table 5**). This bioinformatics suggest that manifestation of HR loss is prominently observed in breast tumors exhibiting the luminal progenitor signature and underscore the critical requirement of HR for genome integrity particularly in this luminal progenitor population.

**Fig. 6.**
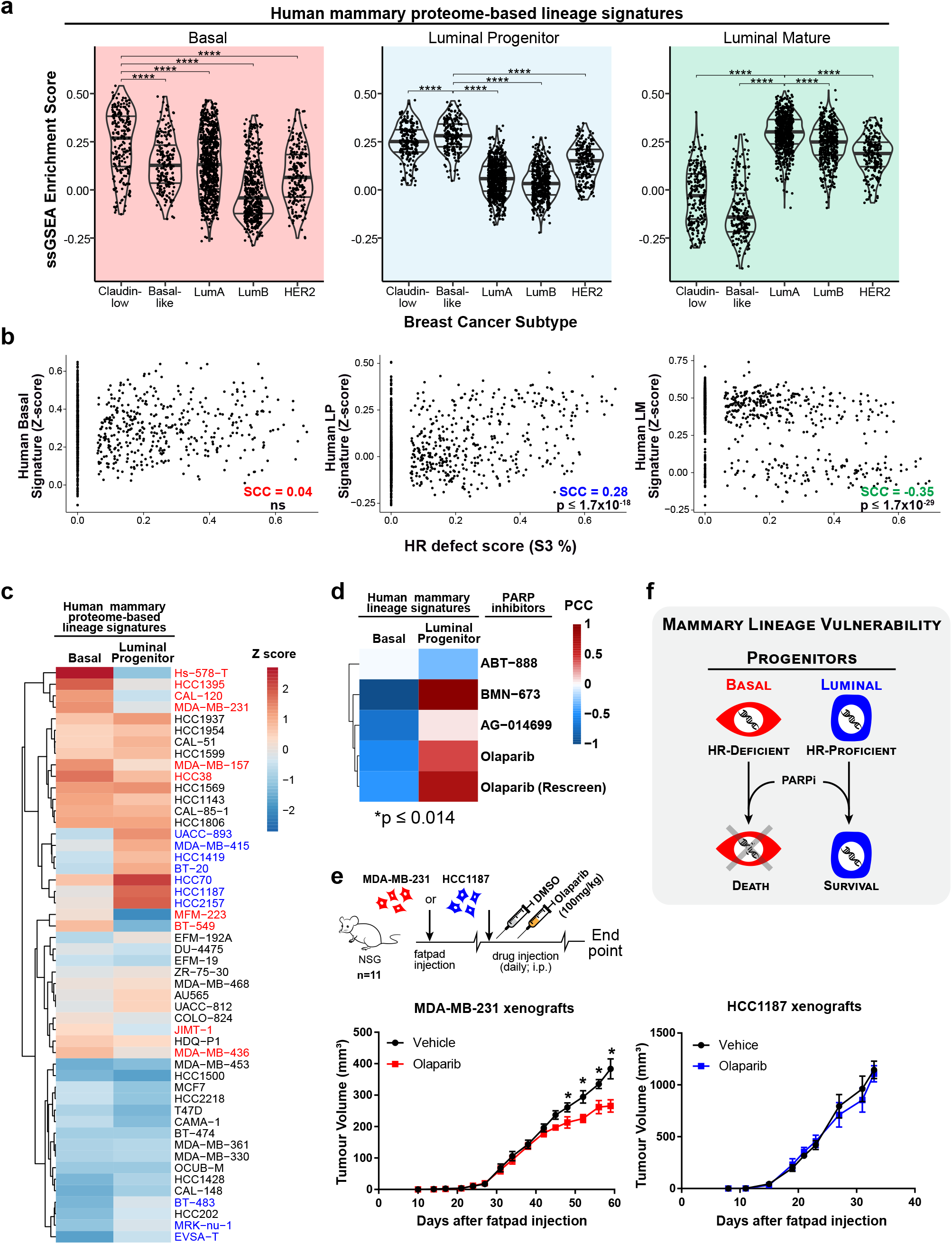
Normal human mammary lineage signatures predict sensitivity to PARPi in human breast cancer cell lines. **(a)** Violin plots comparing our normal human mammary basal, luminal progenitors, and luminal mature lineage signatures (one cell type was compared to the other two; fold-change > 5; p<0.05 from One-way ANOVA in conjunction with Tukey’s test) with mRNA expression for each of the 5 major breast cancer subtypes. P-values were adjusted for multiple testing. **** P<0.0001. **(b)** Scatter plots depicting the distribution of our human mammary lineage signature score and of somatic mutation signature #3, which represents HR defect (inferred percentage contribution in each tumor). The Spearman’s correlation coefficients (SCCs) and their corresponding p-values are indicated. ns, not significant. **(c)** Heatmap showing unsupervised hierarchical clustering of 50 human breast cancer (hBC) cell-line gene expression profiles based on normal human basal and luminal progenitor proteome-defined lineage signatures. The top 10 hBC cell-lines that display the largest differential enrichment (based on the ssGSEA scores) between basal and luminal progenitor signatures are highlighted in red or blue, respectively. **(d)** Heatmap of Pearson correlation coefficient (PCC) values corresponding to the correlations between human mammary lineage signature-breast cancer cell-line ssGSEA scores and IC50 values from GDSC. The results are shown for relevant DNA damage-related agents including 5 PARPi (p≤0.014). The p-value was calculated using unpaired t-test to determine differential PARPi sensitivity of the top 10 human breast cancer cell-lines enriched for basal or luminal progenitor lineage signature. **(e)** Schematic diagram illustrates olaparib dosing of human breast cancer (hBC) xenograft tumours. When the basal signature-enriched MDA-MB-231 or luminal progenitor-enriched HCC1187 tumours had reached 100-200 mm^3^ (n=22 per cell-line), mice were randomized into two groups and were treated with either vehicle control (10% DMSO in saline solution) or 100 mg/kg olaparib in 10% DMSO daily via intraperitoneal injections until they reached an end-point. The growth of olaparib-treated MDA-MB-231 (red) or HCC1187 (blue) tumour xenografts are plotted beside the DMSO control group (black). All data represented as mean ± SEM. P-values were determined by multiple t-test. *q<0.001. **(f)** A model summarizing mammary lineage-dependent HR capacity and PARP inhibitor (PARPi) vulnerability. Normal basal progenitors are selectively targeted by PARPi due to an intrinsic functional HR deficiency. Meanwhile, HR-proficient normal luminal progenitors are able to efficiently repair PARPi-induced DNA double-strand breaks, and therefore are intrinsically resistant to PARPi.

We then compared our basal and luminal progenitor signatures with 50 human breast cancer cell-lines from the Genomics of Drug Sensitivity in Cancer database^49^ that catalogues their drug responses to >100 anti-cancer agents. We selected the top 10 cell-lines that showed enrichment for either basal or luminal progenitor IDs (**Fig. 6c** and **Extended Data Fig. 9a**) and correlated them with their drug sensitivity data to known 23 DNA-damaging agents, including 5 PARPi (**Fig. 6d** and **Supplementary Table 6)**. Notably, the basal and luminal progenitor-enriched cell-lines showed differential responses to the DNA-damaging agents (**Extended Data Fig. 9b**). Specifically, the cell-lines with a strong basal ID had greater sensitivity (i.e. negatively correlated) to the PARPi response than those with a luminal progenitor ID (p≤0.014; **Fig. 6d** and **Supplementary Table 7**). Our lineage IDs also predicted PARPi sensitivity in the order of the magnitude of PARP-trapping potency: the PARPi that have the least or lesser trapping potency, such as ABT-888 (veliparib) and AG-014699 (rucaparib), showed less differential sensitivity compared to olaparib and BMN-673 (talazoparib) which are known to have higher or the highest trapping potency^50^ (**Fig. 6d**). Therefore, normal mammary lineage ID predicted the sensitivity of breast cancer cell lines to PARPi and may be applicable to other chemotherapeutics.

Finally, we examined whether our lineage IDs can predict PARPi response *in vivo*. Four human breast cancer cell lines enriched for basal ID and three enriched for luminal progenitor ID were engrafted into the inguinal mammary gland of 6-7 week-old virgin female NSG mice; only MDA-MB-231 and HCC1187 cell-lines grew as tumour xenografts *in vivo*, representing “basal” and “luminal progenitor” breast tumors, respectively (**Extended Data Fig. 10a**). Both cell-lines are triple-negative for estrogen receptor, progesterone receptor and HER2 status; MDA-MB-231 appears to be mesenchymal-like and has been closely aligned to the Claudin-low tumour subtype^51^, whereas HCC1187 closely resembles the Basal-like tumour subtype^52^. Our flow cytometry analyses of dissociated fully grown xenografts showed that MDA-MB-231 expressed CD44, a mesenchymal stem cell marker, in addition to low expression of human epithelial cell marker (h-EpCAM). In contrast, HCC1187 lacked CD44 and highly expressed EpCAM (**Extended Data Fig. 10b**). We treated xenografts daily with either olaparib (100 mg/kg; i.p.) or vehicle control until the mice reached the endpoint (**Fig. 6e**). “Luminal progenitor” HCC1187 tumors showed no difference in growth upon treatment, while olaparib significantly impeded the growth of “basal” MDA-MB-231 tumors (Multiple t-test; FDR < 0.01; **Fig. 6e**). These data demonstrate that the intrinsic olaparib resistance identified here for the normal luminal lineage also prevails in the context of breast cancer xenografts. Furthermore, tumour selection based on lineage-specific signatures has the potential to identify better responders for achieving greater olaparib efficacy.

## Discussion

Our understanding of DDR has been limited by the dogma that every cell has the same DSB repair capacity. Although others have shown variances in DNA repair within the mammary lineages by examining telomere-associated^53^ or oncogene-mediated DDR^54^ and by using an HR-reporter mouse^55^, cell cycle-dependent DSB end-resection has remained the overarching principle known to dictate the choice between HR and NHEJ^28^. We identify cell lineage as a novel imperative factor in DSB repair pathway choice in the mammary gland; the luminal lineage is HR-proficient, whereas the basal lineage is innately limited in HR capacity and thereby exhibits “BRCAness”^32^. We propose that mammary lineage commitment confers each epithelial lineage with defined DSB repair machinery which is then regulated by the cell cycle.

Current determinants of PARPi sensitivity are based on screening for mutations that confer HR deficiency. However, both experimental data^56^ and clinical trials^57–59^ have documented that a *BRCA* mutation alone does not entirely predict PARPi benefit indicating that factors beyond *BRCA* mutation are involved. In fact, OlympiAD^60^ and EMBRACA^61^ breast cancer trials testing PARPi as single-agent therapy have shown significant improvement, but only by 3 months, in progression-free survival of *BRCA1/2*-mutated breast cancer patients. This limited success may be explained by our finding that luminal progenitors, the putative cell-of-origin for *BRCA1* breast cancers^46,47^, are intrinsically PARPi-resistant, and therefore might not have been effectively eliminated by PARPi monotherapy (**Fig. 6f**). Work presented here underscores the impact of mammary lineage on drug response. As such, identification of a patient subgroup whose tumours exhibit a basal ID and/or reduced luminal progenitor ID may present a method for pinpointing better responders to PARPi therapy, as demonstrated in our xenografts representing triple-negative breast cancers.

To potentiate PARPi efficacy, combination strategies may be useful especially if they can simultaneously deplete more than one type of precursor cell/cell-of-origin. Numerous combination strategies have been tested in *in vivo* experiments and are currently being tested in the clinic, including chemotherapy^62^ (NCT03150576; NCT02163694), targeted therapy^63^ (NCT02208375; NCT01623349), immunotherapy^64^ (NCT02484404, NCT02849496, NCT02734004, NCT03594396, NCT03167619), and endocrine therapy^65,66^. Our work has demonstrated that lineage-specific DSB repair features explain the possible shortcomings of olaparib alone in *BRCA* breast cancer patients and offers an opportunity to improve patient stratification to benefit high-risk women.

## Author Contributions

H.K. designed and performed majority of the experiments and data analyses. A.E.C. generated mouse and human primary mammary proteomes and developed a workflow of mouse and human CFC assays. L.P. and M.A.P. performed bioinformatics on TCGA and interrogated GDSC drug sensitivity dataset. K.A. assisted in vivo drug injections and performed bioinformatics on METABRIC and GDSC drug sensitivity datasets. S.N. and P.D.W. assisted in vivo engraftment. S.H., G.D.B guided GSEA pathway analysis and enrichment map visualization. M.P. operated ImageStream®^X^ Mark II instrument and guided post-acquisition data analyses. M.M. and S.N. assisted in processing primary human breast specimens and human CFC assays. S.H. and H.K.B. acquired human breast specimens. T.K. generated mouse and human global mammary proteomes. H.K., P.D.W., M.M. and R.K. wrote and edited the manuscript. R.K. recruited funding and supervised the study.

## Supplementary Material

### Methods

#### Mice

8-10 week-old virgin female FVB/NJ mice were purchased from the Jackson Laboratory or Charles River Laboratories. 8-10 week-old virgin female *K14-cre*^+^;*Brca1^fl/fl^* transgenic mice^35^ were bred in-house at more than 10 back-crosses in the FVB/NJ background. 12–20 week-old, virgin female B6;129-*Trp53bp1^tm1Jc^*/J mice^67^ were provided by Dr. Daniel Durocher. All FVB female mice were subcutaneously implanted with a 14-day slow-release pellet of 0.14 mg 17β-estradiol plus 14 mg progesterone (E+P; Innovative Research of America) one week after bilateral ovariectomy. These mice were sacrificed on the 14^th^ day of E+P treatment with or without whole-body irradiation. 4 week-old virgin female NSG mice were purchased from MaxBell Basement ARC Barrier Facility for xenograft experiment. All procedures were conducted in compliance with the Canadian Council for Animal Care guidelines under protocols approved by Animal Care Committee of the Princess Margaret Cancer Centre, Toronto, Ontario, Canada.

#### Irradiation

##### In vivo

Mice were exposed to a single dose of 6 Gray (Gy) from a Cs-137 irradiator (Gammacell^®^ 40 Exactor) and sacrificed at a specified time-point after irradiation.

##### In vitro

Primary mouse mammary epithelial cells seeded in a 6-well cell culture plate (Greiner, 657160) were irradiated at a single dose of 3 Gy (Gammacell® 40 Exactor) on day 1 of the CFC assay.

#### Primary human breast tissue

All human breast tissues were obtained from women undergoing prophylactic mastectomies within 24-48 hours of surgery at the Princess Margaret Cancer Centre, Toronto, Canada, under full informed consent and in accordance with Institutional Research Ethics Board approval. Tissues were then processed as described previously^5,68^. Briefly, surgically excised breast tissues were digested with 300 U/mL collagenase (Sigma, C9891) plus 100 U/mL hyaluronidase (Sigma, H3884) to isolate mammary organoids, which were cryopreserved in liquid nitrogen until processed for single-cell suspensions.

#### Primary mammary single-cell suspensions

##### Mouse

Mouse single-cell suspension was performed as described previously^10^. Briefly, freshly harvested mammary glands were minced and incubated in DMEM/F12 media with 300 U/mL collagenase plus 100 U/mL hyaluronidase (STEMCELL Technologies, 07912) at 37°C for 1.5 h. Resulting mammary organoids were serially treated with ammonium chloride (STEMCELL Technologies,07850), 0.25% trypsin-EDTA (STEMCELL Technologies, 07901) and 5 U/mL dispase (STEMCELL Technologies, 07913) plus 50 µg/ml DNase I (Sigma, D4513). The resulting cells were filtered through a 40 µM cell strainer (Fisher Scientific, 22-363-547).

##### Human

Human single-cell suspensions were prepared as reported previously^5,68^. Briefly, previously processed human mammary organoids were thawed from liquid nitrogen and serially digested as above with 0.25% trypsin-EDTA and 5 U/mL dispase and 50 µg/ml DNase I. The resulting cells were filtered through a 40 µm cell strainer.

#### γ-H2AX intracellular flow cytometry

Freshly dissociated mouse mammary single cells were stained with a viability dye (Zombie UV; BioLegend, 423107, 1:100) and a cocktail of cell surface markers to exclude lineage^+^ (Lin^+^), including: PE/Cy7-conjugated antibodies to CD45 (hematopoietic; eBioscience, 25-0451-82, 1:800), CD31 (endothelial; eBioscience, 25-0311-81, 1:200), Ter119 (erythrocyte; eBioscience, 25-5921-82, 1:100) cells. Anti-CD24-PerCP-eFluor^®^ 710 (eBioscience, 46-0242-80, 1:400), anti-CD49f-APC (R&D, FAB13501A, 1:100), anti-CD49b-PE (BioLegend, 103506, 1:250), anti-Ly-6A/E (Sca-1)-APC/Cy7 (BioLegend, 108126, 1:500) were used to segregate basal, LP, LM, and stromal populations. After cell surface staining, the cells were fixed in 4% paraformaldehyde (PFA) for 10 min at room temperature, washed in PBS, and stored overnight at 4°C. The next day, the cells were permeabilized with 0.1% Triton-X/PBS for 5 min at room temperature and stained with Alexa Fluor^®^ 488-conjugated anti-phospho-histone H2A.X (Ser139; γ-H2AX; Cell Signaling Technology, 9719S, 1:200) or concentration matched Alexa Fluor^®^ 488-conjugated rabbit IgG Isotype (Fc) control (Cell Signaling Technology, 4340S). Flow cytometry acquisition was performed using BD LSRFortessa™ and analyzed with FlowJo software (Tree Star, Inc.).

#### Amnis Imaging flow cytometry

##### Staining

Freshly dissociated mouse single cells were stained with a fixable viability Zombie UV Dye (BioLegend, 423107, 1:100) and a cocktail of cell surface markers including: biotin-conjugated CD45 (eBioscience, 13-0451-82, 1:800), CD31 (eBioscience, 13-0311-8, 1:200), Ter119 (eBioscience, 13-5921-81, 1:100) which were subsequently labelled with secondary conjugate streptavidin-eFluor 450™ (eBioscience, 48-4317-82, 1:500); anti-CD24-APC-eFluor^®^ 780 (eBioscience, 47-0242-82, 1:400), anti-CD49f-PE/Cy7 (BioLegend, 313622, 1:100), anti-CD49b-PE (BioLegend, 103506, 1:250), anti-Ly-6A/E(Sca-1)-PE-CF594 (Sca-1; BD Biosciences, 562730, 1:500). After washing, the surface stained cells were fixed in 4% PFA for 10 min at room temperature, washed in PBS, and stored overnight at 4°C. The next day, the cells were permeabilized with 0.1% Triton-X/PBS for 5 min at room temperature, washed and stained with anti-Rad51 (H-92; SantaCruz, sc-8319, 1:50) or anti-DNA-PKcs (phospho S2056; Abcam, ab18192, 1:1,500) which were subsequently labelled with goat anti-rabbit Alexa Fluor^®^ 488-conjugated secondary antibody (Thermo Scientific, A-11008, 1:200). Nuclear DNA was stained by DRAQ5™ (ThermoFisher Science 62251, 2.5 μM). Imaging flow cytometry was performed on an ImageStream®^X^ Mark II (Excitation lasers: 405, 488, 561, 592, 642nm; MilliporeSigma) with INSPIRE^®^ software.

##### Image acquisition

Stained cells were resuspended in a volume of 50 μl PBS + 1% FBS in a 1.5 mL low retention microfuge tube (Sigma, T4816). Samples were then acquired on a 5 laser 12 channel ImageStream®^X^ Mark II imaging flow cytometer at 60X magnification following ASSIST calibration (Amnis^®^ Corporation). Channels 1, 2, 3, 4, 6, 7, 9, 11 and 12 were used for acquisition along with lasers 405 nm (100 mW), 488 nm (150 mW), 561 nm (200 mW), 592 nm (300 mW) and 642 nm (150 mW) for excitation. A bright-field (BF) area lower limit of 50 μm^2^ was used to eliminate debris and calibration beads during sample acquisition, while samples were collected in a series of 50×10^3^ event raw image files (.rif). For single stained compensation controls, BF illumination was turned off and approximately 3000 events within the positive signal fraction were acquired. An initial compensation matrix was generated by loading the single stained raw image files into the IDEAS compensation wizard (IDEAS^®^ version 6.2) with further refinements to the compensation matrix made as necessary through manual adjustment^69,70^. Once generated the compensation matrix was then be applied to the sample raw image files to create compensated image files (.cif) which were then analyzed.

##### High-throughput DSB repair foci counting

Analysis was carried out using the IDEAS^®^ Software (version 6.2, Amnis Corporation). Using masks and features as defined in the IDEAS reference manual (version 6.0, Amnis Coporation). Briefly, the analysis workflow including IDEAS formatted axis feature/mask descriptors in parenthesis was as follows: (a) Gate on focused cells using the gradient RMS feature and default M01 mask in the BF channel (Gradient RMS_M01_Ch01). (b) Gate on single cells by plotting features aspect ratio vs area within the M01 mask in the BF channels (Area_M01 vs. Aspect Ratio_M01). (c) Gate on circular cells by plotting features aspect ratio vs circularity within the M01 mask in the BF channels (Circularity_M01 vs. Aspect Ratio_M01). (d) Gate on the viable lineage negative (Lin^−^) population by plotting the intensity feature of the viability + lineage negative stain vs. area (both viability dye and lineage negative panel are detected in the same channel (Intensity_MC_Ch07 Lin neg/viability vs. Area_M01). Lin^−^ gate was determined by using fluorescence minus one (FMO) control. (e) The stromal, luminal and basal populations were distinguished by plotting the intensity features of CD49f vs. CD24 (Intensity_MC_Ch06_CD49f vs. Intensity_MC_Ch12_CD24). (f) Further subdivision of the luminal population into luminal progenitor (LP) and luminal mature (LM) populations was achieved by plotting the intensity features of CD49b vs. intensity of Sca1 (Intensity_MC_Ch03_CD49b vs. Intensity_MC_Ch04_Sca1). (g) The number of RAD51 or pDNA-PKcs foci in basal, LP, and LM populations were then quantified using the “Spot Count” feature based on three different user-defined masks that define nucleus by DRAQ5 signal and repair foci by signal/intensity and size. The following feature/mask parameters were used to count Rad51 or pDNA-PKcs foci in single cells. Rad51: Spot Count_Morphology(M11, Ch11) And Intensity(M02, Ch02, 1200-4095) And Spot(M02, Ch02, Bright, 9.5, 1, 0)_4. pDNA-PKcs: Spot Count_Morphology(M11, Ch11) And Intensity(M02, Ch02, 1000-4095) And Spot(M02, Ch02, Bright, 9.5, 1, 0)_4

#### Fluorescence Activated Cell Sorting (FACS) staining

Freshly dissociated mouse mammary epithelial cells were stained with the following antibodies: PE/Cy7-conjugated lineage antibodies as described above, and various combinations of anti-CD24-APC-eFluor^®^ 780 (eBioscience, 47-0242-82, 1:400), anti-CD24-PerCP-eFluor^®^ 710 (eBioscience, 46-0242-80, 1:400), or anti-CD24-FITC (BD Biosciences, 553261, 1:400), and anti-CD49f-APC (R&D, FAB13501A, 1:100) or anti-CD49f-FITC (BioLegend, 313606, 1:100). Dead cells were excluded with DAPI (5 mg/mL; 1:10,000). Cell sorting was performed on a BD FACSAria™ II.

#### Human mammary epithelial flow cytometry

Freshly dissociated human mammary epithelial cells were stained with antibodies against anti-CD45-PE/Cy7 (BioLegend, 304016, 1:200), anti-CD31-PE/Cy7 (BioLegend, 303118, 1:50), anti-CD326 (EpCAM)-PE (BioLegend, 324206, 1:50) and anti-CD49f-FITC (BioLegend, 313606, 1:100). Dead cells were excluded with Zombie UV Dye (BioLegend, 423107, 1:100). Flow cytometry analysis was performed by using BD™ LSR II or BD LSRFortessa™ and FlowJo software (Tree Star, Inc.).

#### *In situ* immunofluorescence staining

Freshly harvested mouse mammary glands were fixed in 4% PFA at 4°C overnight and stored in 70% EtOH prior to dehydration and embedding into paraffin blocks. Tissue-sections were deparaffinized and rehydrated prior to antigen retrieval in a pressure cooker (BioMedical) for 30 minutes at 121°C in pH 6.0 citrate buffer. The sections were blocked with 5% goat serum, 1% glycerol, 0.1% Bovine Serum Albumin, 0.1% cold water fish skin gelatin (Sigma, G7765) in PBS for 1 h at room temperature. The sections were stained with a cocktail of primary antibodies that were diluted in the same blocking buffer overnight at 4°C. Next day, the tissue sections were washed in 0.1% Tween-20/PBS and then incubated with secondary antibodies diluted in blocking buffer for 1 h at room temperature. The sections were mounted in Anti-fade ProLong^TM^ Gold with DAPI (Thermo Scientific, P36935). Primary antibodies used are as follows: anti-Keratin 14 (K14; BioLegend, 906004, 1:400), anti-Ki67 (BioLegend, 652402, 1:400), anti-phospho-Histone H2A.X (Ser139; Cell Signaling Technology, 9718, 1:200), anti-Rad51 (SantaCruz, sc-8319, 1:50), anti-DNA-PKcs (phospho S2056; Abcam, ab18192, 1:400), anti-progesterone receptor (PR; LifeSpan Biosciences Inc., LS-B5236, 1:400), anti-PARP1 (LifeSpan Biosciences Inc., LS-B3432, clone A6.4.12, 1:100), anti-PAR (Cell Signaling Technology, #83732, 1:100). Secondary antibodies used are as follows: anti-rat conjugated to AlexaFluor^®^ 647 (Thermo Scientific, A-21247), anti-chicken-Cy™3 AffiniPure (Jackson ImmunoResearch, 103-165-155), and anti-rabbit-AlexaFluor^®^ 488 (Thermo Scientific, A11008).

#### Confocal microscopy image acquisition

All tissue immunofluorescence images were acquired on a Zeiss LSM 700 confocal microscope using a 40X oil-immersion objective lens. At least 3 or 4 Z-planes were acquired.

#### *In situ* foci counting

Z-projected confocal images were analyzed to count nuclear foci in ImageJ, as described previously^71^. Briefly, DAPI and Rad51 channels were used to identify all nuclei and foci in the image, respectively. Based on the K14 channel, each nucleus was annotated as “basal” (K14^+^), “luminal” (K14^−^ cells surrounded by K14^+^ cells) or “stromal” (K14^−^ cells surrounding K14^+^ cells). R software was used to count the total number of nuclei displaying ≥ 2 foci (i.e. Rad51^+^ cells) in each population.

#### Colony forming cell (CFC) assay

##### Mouse

Mammary 2D colony-forming assays were performed as described previously^5^. FACS-purified mouse total basal and luminal cells were plated in 6-well plate (Greiner, 657160) with 2 x 10^5^ irradiated NIH-3T3 feeder cells per well in Mouse EpiCult media (STEMCELL Technologies, 05610) supplemented with 5% FBS, 20 ng/mL hFGF (STEMCELL Technologies, 02634), 10 ng/mL bovine EGF (STEMCELLTechnologies, 78006), 4 μg/mL heparin (STEMCELL Technologies, 07980), 5 μM ROCKi (Millipore, SCM075), Antibiotic-Antimycotic Solution (Wisent, 450-115-EL; 1:100) and incubated in a low oxygen (5%) incubator at 37°C. On the 7^th^ day, colonies were fixed with 1:1 acetone:methanol (v/v) and stained with Giemsa (Fisher, 264-983) according to their manufactor’s protocol. The total number of colonies were manually counted under a Leitz dissecting microscope.

##### Human

Breast 2D colony-forming assays were performed with patient samples as described previously^5,36,68^. Total dissociated breast cells were seeded with irradiated NIH 3T3 cells in 60 mm culture dishes (Greiner, 82050-546) that were collagen-coated for 1 h (STEMCELL Technologies, 04902). Single-cell suspensions of freshly dissociated human breast cells were cultured in Human EpiCult-B (STEMCELL Technologies, 05601) at 5% oxygen for 10 days. On day 1, the media was changed to serum (FBS)-free conditions till endpoint. On the 10^th^ day, colonies were fixed, stained and counted as above.

#### Small molecule inhibitors/*In vitro* drug testing

All drugs were dissolved in DMSO such that the final concentration of DMSO did not exceed 0.1% (v/v) in each well/dish. Small molecule inhibitors used were: olaparib (AZD-2281; MedChem Express, HY-10162; Selleck Chemicals, S1060), talazoparib (BMN-673; MedChem Express, HY-16106), and KU-57788 (NU7441; Selleck Chemicals, HY-11006). For *in vitro* drug treatment, the drugs were added into each well or the dishes on day 1 of all CFC assays.

#### Neutral comet assay

FACS-sorted basal, LP and LM cells were resuspended in PBS at 20,000 cells/mL and were mixed with 0.7% low-melting point agarose gel at 1:10 ratio. Each population mixture was laid on a glass slide pre-coated with 1% normal melting point agarose and covered with an 18×18 mm coverslip. The coverslips were carefully removed after gel solidification at 4°C, and the slides were then processed according to the manufacturer’s recommendation (Trevigen Neutral CometAssay®). Briefly, slides were immersed in Lysis solution (Trevigen; 4250-050-01) at 4°C overnight and then immersed in 1X Neutral Electrophoresis buffer for 30 min. The slides were transferred to the CometAssay® electrophoresis unit filled with chilled 1X Neutral Electrophoresis buffer and electrophoresed at 21 V for 7 min at 4°C. The slides were then fixed in DNA Precipitation Solution followed by 70% ethanol for 30 min each at room temperature. The slides were dried at room temperature and stained with SYBR® Gold gel staining solution (Thermo Fisher, S-11494, 1: 10,000). The Olive tail moment was measured by using Komet Software (Andor Technology), and at least 100 comets were analysed per population per mouse.

#### Gene set enrichment analysis (GSEA)

Protein expression profiles^5^ of basal, LP, and LM populations from adult female mice that were treated with estrogen plus progesterone were included in the pathway analysis by using GSEA, and those exposed to estrogen alone were excluded in this analysis. GSEA was performed as described previously^23^. Briefly, protein expression profiles of each mammary population were compared to those of the other two populations (“Rest”) to determine biological pathways that were uniquely upregulated or downregulated in that population. The Gene Matrix Transposed (.gmt) or Mouse_GOBP_AllPathways_no_GO_iea_August_01_2017_UniProt.gmt file was accessed from the Bader Lab gene sets website (http://download.baderlab.org/EM_Genesets/current_release/) and used for GSEA and enrichment map visualization. GSEA parameters were defined as suggested by the Bader Lab GSEA Tutorial website (https://enrichmentmap.readthedocs.io/en/docs-2.2/Tutorial_GSEA.html#step-1-generate-gsea-output-files).

#### Enrichment map visualization

GSEA results were visualized by using the Enrichment Map App in Cytoscape v3.5.1. Only the upregulated gene sets or pathways that have met the cut-off of FDR < 0.01 were clustered by using the MCL cluster algorithm in the clusterMaker App. Then, each cluster was manually annotated with a common biological theme. A node (circle) represents a single pathway defined by publicly available databases curated by the Bader Lab. An edge (line) represents the extent of shared genes between the two nodes.

#### Heatmap of mouse DDR protein abundance

Protein abundance values of the genes curated in the “DNA double-strand break repair” pathway (R-HSA-5693532; Reactome Database ID Release 63) was queried in our previously published mouse mammary proteomic dataset^5^. The heat map shows the z-scores of LFQ-adjusted IBAQ values in basal, luminal progenitor (LP), and luminal mature (LM) cell types from estrogen plus progesterone (E+P) samples.

#### Human breast cancer cell-lines

All 7 human breast cancer cell-lines used in this study were purchased from either ATCC or DSMZ. MDA-MB-231 (ATCC^®^ HTB-26™), Hs 578T (ATCC^®^ HTB-126™), EFM-192A (ACC 258), and EVSA-T (ACC 433) were cultured in DMEM media + 10% FBS. HCC1395 (ATCC^®^ CRL-2324™), BT-549 (ATCC^®^ HTB-122™), and HCC1187 (ATCC^®^ CRL-2322™) were cultured in RPMI media + 10% FBS. All cell-lines were incubated at 37°C, 5% CO_2_.

#### Human breast cancer cell-line engraftment

Each human breast cancer cell-line in PBS was mixed with Matrigel® (Corning, 356231) in 1:1 v/v ratio. A total of 10 μl cell-Matrigel mixture containing 5×10^5^ cells was injected directly into the right inguinal mammary gland of 6-7-week-old virgin female NSG mice using a Hamilton syringe. Xenograft tumours were monitored twice a week starting 7 days after engraftment. Tumour dimensions were measured with a Vernier caliper two times a week and tumour volume (mm^3^) was estimated by 0.5 × (minimum diameter in mm)^2^ × (maximum diameter in mm) from day 7 post-injection until the end of the study. Mice were sacrificed when humane endpoints were reached (tumour volume >1500 mm^3^, cumulative clinical score >8 or a max score for any animal condition, or limb paralysis).

#### *In vivo* drug testing

1 g of olaparib (MedChemExpress, HY-10162) was dissolved in 10 mL of DMSO and stored at - 80°C. A fresh aliquot of 100 mg/mL olaparib stock was diluted 1:10 in the vehicle solution (10% w/v 2-hydroxypropyl-β-cyclodextrin in PBS) at the time of injection. Once MDA-MD-231 and HCC1187 tumors reached a volume of 100-200 mm^3^, mice were randomized into two groups and were treated with either vehicle control (10% DMSO in the vehicle solution) or 100 mg/kg olaparib daily via intraperitoneal injections until they reached an endpoint as described above.

#### Human DDR proteomic analysis

The expressions of DDR proteins in the three mammary epithelial populations: basal (BC), luminal progenitor (LP), and luminal mature (LM) was extracted from our previously published human mammary epithelial proteome^6^. The proteomic dataset contains log2-transformed iBAQ-adjusted LFQ values representing protein abundance which were adjusted for batch effects using the ComBat function in the ‘sva’ R package (version 3.30.1). Only samples from premenopausal patients were taken into account (n=6 for each BC, LP, LM cell types). Of the 276 curated human DDR genes^39^, 127 proteins were detected in the mammary proteomic dataset based on matching by gene symbol.

#### Volcano plot and Pathway analysis on the human DDR proteins

Of 127 DDR proteins, proteins were defined as highly expressed or enriched in one cell type if they met the fold-change (FC) and statistical significance cut-offs compared to the other two cell types. Enriched proteins had a log2FC > 0 and a p-value < 0.05 (paired t-test; p-value was adjusted for multiple testing via FDR) and were visualized in a volcano plot. The top 5 “upregulated” and “downregulated” proteins were based on rank values from the summation of p-value and fold-change ranks. The protein with the lowest p-value was given a rank of 1, while the highest earned the rank of r, which represents the number of total proteins (i.e. 127). For upregulated, the protein with the most positive log2FC value was given a rank of 1, while the least positive (log2FC > 0) value was given a rank of r. For downregulated, the protein with the most negative log2FC value was given a rank of 1, while the least negative (log2FC < 0) value was given a rank of r. The gene names of all enriched proteins were visually labelled in volcano plots. Pathway analysis of all enriched proteins/genes for each cell type queried KEGG 2019 terms using Enrichr (https://amp.pharm.mssm.edu/Enrichr/). The top 5 enriched pathways was determined based on the ‘combined’ score and were visualized in bar charts.

#### Generation of human and mouse lineage signatures

The total mammary proteome consisted of 6034 (human) and 4695 (mouse) annotated proteins. Basal, luminal progenitor and luminal mature lineage signatures were acquired by searching for proteins enriched in one cell type compared to the other two cell types with a fold-change > 5 in human or fold-change > 3 in mouse and p < 0.05 in one-way ANOVA in conjunction with Tukey’s test.

#### Correlations to breast cancer subtypes

Human and mouse lineage signatures in breast cancer expression profiles from METABRIC via single-sample Gene Set Expression Analysis (ssGSEA) using the ‘GSVA’ R package (version 1.30.0). The scores were categorized by PAM50 plus Claudin-low subtypes for each signature and assessed for significance using a one-way ANOVA and Tukey’s multiple comparisons test.

TCGA breast cancer RNA-seq and somatic mutation data were obtained following approval by the Data Access Committee (project #11689). The results published here are partly based upon data generated by TCGA managed by the NCI and NHGRI at http://cancergenome.nih.gov.

#### Correlations to COSMIC somatic mutational signatures

Thirty different somatic mutational signatures from COSMIC (Catalogue Of Somatic Mutations In Cancer) were defined using the mutSignatures package in R^48^. The signature scores were computed using the ssGSEA algorithm^72^ with standard parameters and using all genes included in each set. Spearman’s correlation coefficients and p-values were computed in R.

#### Enrichment in human breast cancer cell-lines

Enrichment of human or mouse lineage signatures in 50 human breast cancer cell-lines from the Genomics of Drug Sensitivity in Cancer database was determined using the ssGSEA algorithm. For mouse signatures, gene symbols were converted to human homologs using the ‘biomaRt’ R package (version 2.38.0). Top cell-lines were defined as having the greatest differences between basal and luminal progenitor signatures in ssGSEA enrichment scores.

#### Correlation to breast cancer cell-line drug sensitivity screening

Pearson correlations were performed to measure the association between enrichment scores for each cell type and IC50 values of 23 DDR-related drugs (targeting ‘DNA replication’ or ‘Genome integrity’) in human breast cancer cell-lines from the Genomics of Drug Sensitivity in Cancer portal^49^. Only the top ten cell-lines that were enriched for each basal and luminal progenitor signature were considered. Pearson’s correlation coefficients and p-values were computed in R. The IC50 data was from: https://www.cancerrxgene.org/gdsc1000/GDSC1000_WebResources/Home.html

---

**Extended Data Fig. 1. Optimization of *in situ* DDR detection in the mammary gland after whole-body irradiation.**

**(a)** Immunofluorescence images showing the dose response of γ-H2AX foci formation (green) in the nucleus (DAPI; blue) in the mammary gland of adult virgin female mice 1 h after receiving 2, 4, or 6 Gray (Gy) of irradiation *in vivo*.

**(b)** Time course of 53BP1 (tumor protein p53-binding protein 1; green) foci formation in basal (K14; red) and luminal (K14^−^) cells with a proliferation (Ki67; magenta) marker in the mammary gland harvested at 0.5, 1, 3, 6, or 24 h after 6 Gy *in vivo* irradiation.

**Extended Data Fig. 2. Comprehensive characterization of γ-H2AX *in situ* and by intracellular flow cytometry.**

**(a)** Flow cytometry gating strategy for the exclusion of doublets, dead cells (Zombie UV^+^), and the lineage (Lin^+^) population consisting of hematopoietic (CD45^+^), endothelial (CD31^+^), and erythrocytes (Ter119^+^) from total primary single-cell suspensions from digested mammary gland. The resulting live Lin^−^ cells were separated into basal (red), luminal (black), or stromal (grey) populations based on CD24 and CD49f cell surface markers. The total luminal cells were further separated based on Sca1 and CD49b cell surface markers into luminal progenitor (LP; blue) and luminal mature (LM; green) populations.

**(b)** Time course showing the proportion of γ-H2AX^+^ cells in mammary epithelial and stromal populations after *in vivo* irradiation (6 Gy). One-way ANOVA with Dunnett’s multiple comparison test was applied (n=4-10 mice per time-point). *P<0.05, **P<0.01, ****P<0.0001.

**(c)** Time course showing the median fluorescence intensity (MFI) of stromal γ-H2AX^+^ cells normalized by cell-size (FSC-A; see Extended Data Fig. 2d,e) and the absolute number of γ-H2AX^+^ cells at different time-points after irradiation. One-way ANOVA with Dunnett’s multiple comparison test was applied (n=4-10 mice per time-point). *P<0.05, ***P<0.001.

**(d)** Scatter plot demonstrating median cell-size (FSC-A) of basal, LP, LM, and stromal populations from individual mice (n=70 mice per population). P-values were calculated by one-way ANOVA with Tukey’s multiple comparison test. *P<0.05, ***P<0.001. ns, not significant.

**(e)** Density plot of nuclear area of single-cell images from the ImageStream®^X^ analyses. Nucleus was determined by DRAQ5 stains and the nuclear area calculated by built-in Area function in the IDEAS™ software for basal (n=41,060 cells), LP (n=70,849), LM (n=17,368) and stromal (n = 178,403) populations from a pool of 10 mice. All data represented as mean ± SEM. P-values were determined by ordinary one-way ANOVA. ****P<0.0001.

**(f)** Post-irradiation time course of γ-H2AX (green) foci formation in mammary tissue sections after irradiation (6 Gy; n=4-10 mice per time-point). Basal cells are marked by K14 (red) and proliferating cells by Ki67 (magenta).

**Extended Data Fig. 3. ImageStream®^X^ gating strategies, pDNA-PKcs foci visualization, and absolute counts of Rad51 and pDNA-PKcs-foci containing cells.**

**(a)** A workflow depicting the gating strategy for post-acquisition image processing using the IDEAS™ software. Of all acquired images from ImageStream®^X^ Mark II imaging flow cytometry, focused cells were selected to further gate for single cells defined by morphological features (i.e. aspect ratio, area, and circularity) based on brightfield (CH01) images. The resulting live (Zombie UV^−^), Lin^−^ (lineage (CD45^+^CD31^+^Ter119^+^)-depleted) population was separated based on CD24, CD49f, Sca1, and CD49b cell surface markers to acquire basal, luminal progenitor (LP), and luminal mature (LM) populations.

**(b)** A panel of representative ImageStream®^X^ images displaying pDNA-PKcs (phospho-S2056) foci in the basal, LP, LM, and stromal cells at 3 h post-irradiation. The number of pDNA-PKcs foci was determined using built-in masking techniques that identify nuclear punctate signals based on the DRAQ5 nuclear mask and pDNA-PKcs foci stains. Automatically counted foci number is indicated in yellow at the upper right corner of the ‘Foci Mask’ column.

**(c,d)**, Absolute cell counts displaying 0, 1-4, 5-9, or ≥10 foci of (c) Rad51 or (d) pDNA-PKcs in the basal, LP, and LM populations at un-irradiated (No IR) or 3 h post-irradiation. Nuclear foci were counted in 1,000-7,000 (up to 10,000) mammary epithelial cells per mouse; a total of n=3 mice per treatment group. P-values were determined by one-way ANOVA with Tukey’s multiple comparison test on Rad51^+^ or pDNA-PKcs^+^ cells (i.e. foci number > 0) across all 3 populations. Data represent mean ± SEM ***P<0.001, ****P<0.0001. ns, not significant.

**Extended Data Fig. 4. Characterizing the relationships between Rad51 foci formation and cell proliferation or progesterone receptor status.**

**(a)** Z-projected immunofluorescence imaging of the time course of Rad51 (green) foci formation in mammary glands taken from un-irradiated female virgin mice or post-irradiation (6 Gy) at 0.5, 1, 3, 6, or 24 h. Basal cells (K14; red), and proliferating cells with Ki67 (cyan). Filled and hollow arrowheads indicate Rad51^+^ and Rad51^−^ proliferating (Ki67^+^) cells, respectively.

**(b)** Z-projected immunofluorescence images of Rad51 (green), K14 (red), and progesterone receptor (PR; magenta) in mammary gland at 3 or 6 h post-irradiation *in vivo*.

**(c)** Immunofluorescence images of Ki67 (green) and PR (red) in un-irradiated mammary glands.

**Extended Data Fig. 5. Individual characterizations of primary human breast specimens.**

**(a)** Individual flow cytometric profiles of dissociated single-cells from *BRCA* wild-type, *BRCA1* and *BRCA2*-mutated human primary breast tissue specimens. Each patient specimen was assigned a Roman numeral. Density plots show basal (red; Lin^−^EpCAM^lo^CD49f^hi^), luminal progenitor (LP; blue; Lin^−^EpCAM^hi^CD49f^hi^), and luminal mature (LM; black; Lin^−^ EpCAM^hi^CD49f^lo^) populations separated by EpCAM and CD49f surface markers from live (DAPI^−^), Lin^−^ (CD45^+^CD31^+^-depleted) cells. Corresponding pie charts illustrate the proportion of basal, LP, and LM populations relative to total mammary epithelial cells.

**(b)** Stacked bar charts illustrate relative colony growth in the presence of DMSO vehicle control (D) or olaparib (Ol) of myoepithelial (yellow), bipotent (maroon), and luminal (blue) colonies for individual patient specimens.

**(c)** Age distribution of *BRCA* wild-type, *BRCA1*, and *BRCA2*-mutated patient cohorts (n=5, 9, 6, respectively). P-values were determined by one-way ANOVA with Tukey’s multiple comparison test. ns, not significant.

**Extended Data Fig. 6. Characterizing human DDR proteome of three major mammary epithelial cell types.**

**(a)** Heatmap showing unsupervised hierarchical clustering of detected 127 DDR protein abundance from FACS-purified, normal human basal (red), luminal progenitor (blue; LP), and luminal mature (green; LM) cell populations from premenopausal women who underwent reduction mammoplasty. Normal breast tissues were staged as either follicular (progesterone-low; n=3) or luteal (progesterone-high; n=3) hormonal state. Each DDR gene name is annotated in one or more DDR pathway as shown on the left; Base Excision Repair (BER), Nucleotide Excision Repair (NER), Mismatch Repair (MMR), Fanconi Anemia (FA), Homology-dependent recombination (HDR), Non-homologous End Joining (NHEJ), Direct Repair (DR), Translesion Synthesis (TLS), Nucleotide pools (NP), and Others. Asterisks indicate gene names that have been annotated as a ‘core’ member in a single DDR pathway as defined by Knijnenburg *et al*^39^.

**(b)** Volcano plots showing differential expression of 127 DDR proteins from the human mammary proteomic dataset generated from FACS-purified basal, luminal progenitor (blue; LP) and luminal mature (green, LM) cell populations. The x-axis indicates the relative difference in protein abundance between the two selected populations (log2 fold-change); the y-axis indicates p-values adjusted for multiple correction testing. The grey horizontal line depicts a q-value of 0.05. The gene names of the top 5 significantly upregulated proteins were coloured according to the cell type that were found in for each comparison. Asterisks identify ‘core members’ of one of the 10 DDR pathways defined by Knijnenburg *et al*^39^.

**(c)** Protein abundance for each Parp/PARP member, expressed relative to the highest value within the three cell types. P-values were determined by two-way ANOVA with Tukey’s multiple comparisons test. *P<0.05, **P<0.01, ***P<0.001, ****P<0.0001. ns, not significant.

**Extended Data Fig. 7. Human mammary proteome-based lineage signatures associate with breast cancer subtypes similar to proposed cell-of-origin model.**

**(a, b)** Unsupervised hierarchical clustering of (a) METABRIC or (b) TCGA breast cancer mRNA gene expression profiles based on our human proteome-defined mammary lineage signatures representing basal, luminal progenitor, and luminal mature mammary populations (fold-change>5, p<0.05; One-way ANOVA in conjunction with Tukey’s test compared to the other two cell types). (b) Boxplots depict the expression score of each human lineage signature versus the 4 TCGA breast cancer subtypes.

**Extended Data Fig. 8. Mouse mammary proteome-based lineage signatures associate with breast cancer subtypes similar to proposed cell-of-origin model.**

**(a,b)** Unsupervised hierarchical clustering of (a) METABRIC or (b) TCGA breast cancer mRNA gene expression profiles based on mouse proteome-defined mammary lineage signatures representing basal, luminal progenitor, and luminal mature mammary populations (fold-change > 3, p<0.05 by one-way ANOVA in conjunction with Tukey’s test compared to the other two cell types; see Supplementary Table 4). (a) Violin plots depict the expression score of each of three mouse lineage signatures across 5 breast cancer subtypes. Unpaired t-test was used, and p-values were adjusted for multiple testing. *P<0.05, ***P<0.001, ****P<0.0001. ns, not significant. (b) Boxplots depict the expression score of each mouse lineage signature versus the 4 TCGA breast cancer subtypes.

**(a) (c)** Scatter plots depicting the distribution of normal mouse mammary proteome-defined basal, luminal progenitor, or luminal mature signature (fold-change > 3; p<0.05) scores versus the somatic mutation signature #3, which represents HR defect (inferred percentage contribution in each tumor). The Spearman’s correlation coefficients (SCCs) and their corresponding p-values are indicated. ns, not significant.

**Extended Data Fig. 9. Correlations between human mammary lineage signatures and drug sensitivity to 23 DNA-damaging anti-cancer agents.**

**(a)** Heatmap showing unsupervised hierarchical clustering of 50 human breast cancer (hBC) cell-line gene expression profiles based on our mouse basal and luminal progenitor proteomic signatures. The top 5 hBC cell-lines that display the largest differential enrichment (based on the ssGSEA scores) between basal and luminal progenitor signatures are highlighted in red or blue, respectively.

**(b)** Heatmap of Pearson correlation coefficient (PCC) values corresponding to the correlations between human basal and luminal progenitor signature scores and drug sensitivity (based on normalized half-maximal inhibitory values or IC50) from GDSC^49^. The results are shown for 23 relevant DDR-related agents categorized under ‘DNA replication’ or ‘Genome integrity’. The 5 PARP inhibitors have been bolded.

**Extended Data Fig. 10. *In vivo* and *in vitro* characterization of the 7 human breast cancer cell-lines.**

**(a)** Growth curve of tumour xenografts from 4 basal (red) and 3 luminal progenitor (blue) signature-enriched human breast cancer (hBC) cell-lines in female NSG mice (n = 3-4 per cell-line). Tumour length (mm) was measured twice a week until the end-point. Representative images of fully-grown MDA-MD-231 and HCC1187 xenografts are shown on the bottom. All data represented as mean ± SEM.

**(b)** Flow cytometric analysis on single-cell suspensions from freshly dissociated full size MDA-MD-231 or HCC1187 xenograft tumours. Doublet-excluded, Lin^−^ (CD45^+^CD31^+^-depleted), live (Zombie UV^−^) cells were separated by the human-specific epithelial marker h-EpCAM and the mesenchymal marker CD44.

**Supplementary Table 1: Full report of gene set enrichment analysis (GSEA).**

Detailed ranked lists of pathways found in mouse basal, luminal progenitor (LP) or luminal mature (LM) population compared to the other two populations (“Rest”) based on FACS-purified population-specific protein expression profiles. Gene sets or pathways that have FDR < 0.01 are highlighted in yellow and were used to create the enrichment map for Fig. 1b,c.

**Supplementary Table 2: Complete lists of human KEGG pathway enrichment analyses.**

By using Enrichr^40,41^, pathway analyses was conducted on the human DDR proteins that were found to be significantly upregulated in populations from individual pairwise comparisons as described in Extended Data Fig. 6b. Pathways are ranked in the order of ‘combined’ score.

**Supplementary Table 3: Detailed human gene lists representing the three proteome-based human mammary lineage signatures.**

Three unique groups of gene names that represent proteins upregulated in each of the human basal (Lin^−^EpCAM^lo^CD49f^hi^), luminal progenitor (LP; Lin^−^EpCAM^hi^CD49f^hi^), or luminal mature (LM; Lin^−^EpCAM^hi^CD49f^lo^) populations. Each human mammary lineage signature was acquired by selecting gene names whose protein abundance was over-expressed > 5-fold in one mammary epithelial population compared to the other two (p<0.05; One-way ANOVA with Tukey’s multiple comparison test) based on the iBAQ-adjusted LFQ protein expression values from normal human mammary proteomic dataset^6^.

**Supplementary Table 4. Detailed mouse gene lists representing the three proteome-based mouse mammary lineage signatures.**

Three unique groups of mouse gene names that represent proteins upregulated in each of the mouse basal (Lin^−^CD24^+^CD49f^hi^), luminal progenitor (LP; CD24^+^CD49f^lo^Sca1^−^CD49b^+^), or luminal mature (LM; CD24^+^CD49f^lo^Sca1^+^CD49b^+/−^) populations. Each mouse mammary lineage signature was acquired by selecting gene names whose protein abundance was over-expressed > 3-fold in one mammary epithelial population compared to the other two (p<0.05; One-way ANOVA with Tukey’s multiple comparison test) based on the iBAQ-adjusted LFQ protein expression values from normal mouse mammary proteomic dataset^5^.

**Supplementary Table 5. Complete list of correlations between COSMIC somatic mutation signatures and our human and mouse mammary lineage signatures.**

Full list of Spearman’s rank correlation coefficients and their corresponding p-values were determined between COSMIC’s 30 somatic mutation signatures and our 3 human or mouse mammary lineage signatures.

**Supplementary Table 6. Correlations between human breast cancer cell-lines and our human and mouse mammary lineage signatures.**

Gene expressions of 50 human breast cancer (hBC) cell-lines from GDSC were correlated to our human or mouse proteome-based, basal (BC) and luminal progenitor (LP) lineage signatures. Each hBC cell-line was scored based on the correlations between each signature via single-sample Gene Set Expression Analysis (ssGSEA). Then, score difference (BC.ssGSEA minus LP.ssGSEA) was calculated and sorted in the descending order. The highest score difference correspond to hBC cell-lines that are enriched with the basal signature; the lowest score difference correspond to hBC cell-lines that are enriched with the luminal progenitor signature.

**Supplementary Table 7. Correlations between known drug sensitivity of the human breast cancer cell-lines and our human mammary lineage signatures.**

Pearson’s correlation coefficient was determined for the known drug sensitivity (IC50) of 23 DDR-related agents in the top 10 human breast cancer cell-lines enriched for either basal or luminal progenitor signatures (as determined in Supplementary Table 5).

