## Supplementary Figures for "Mammary lineage dictates homologous recombination repair and PARP inhibitor vulnerability"

Kim et al. Extended Data Fig. 1

a

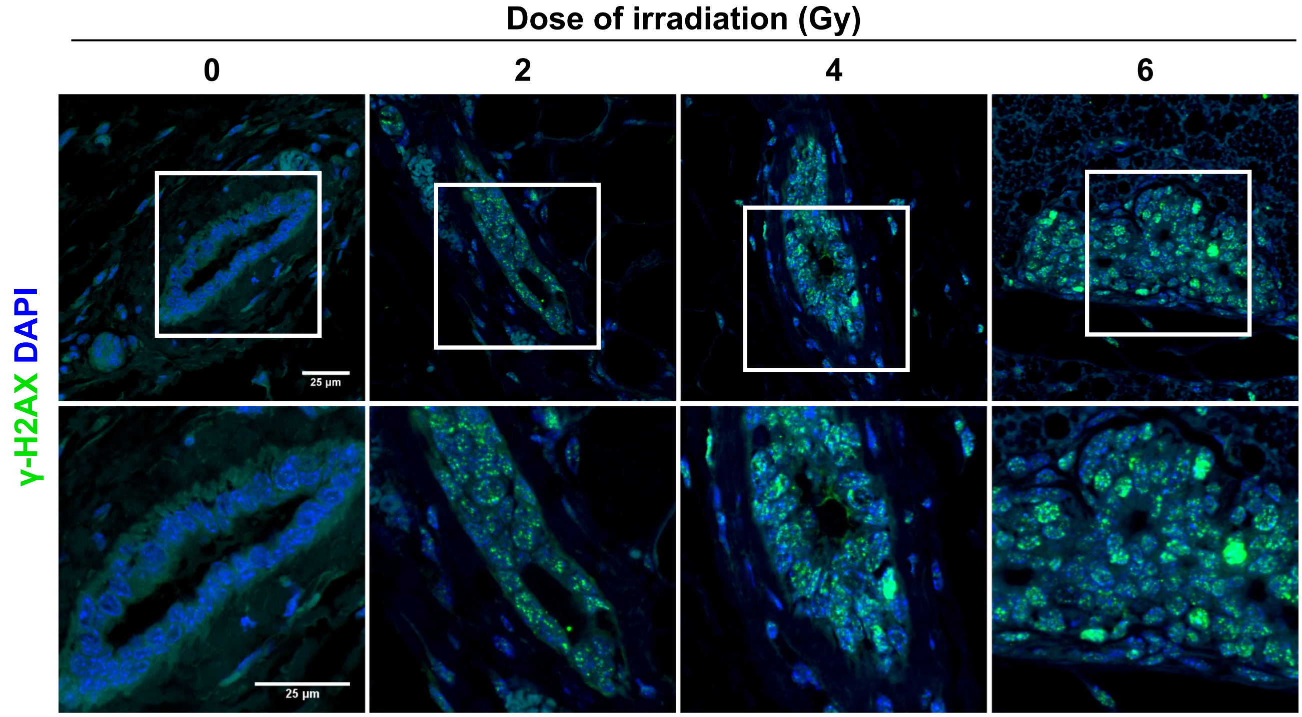


b

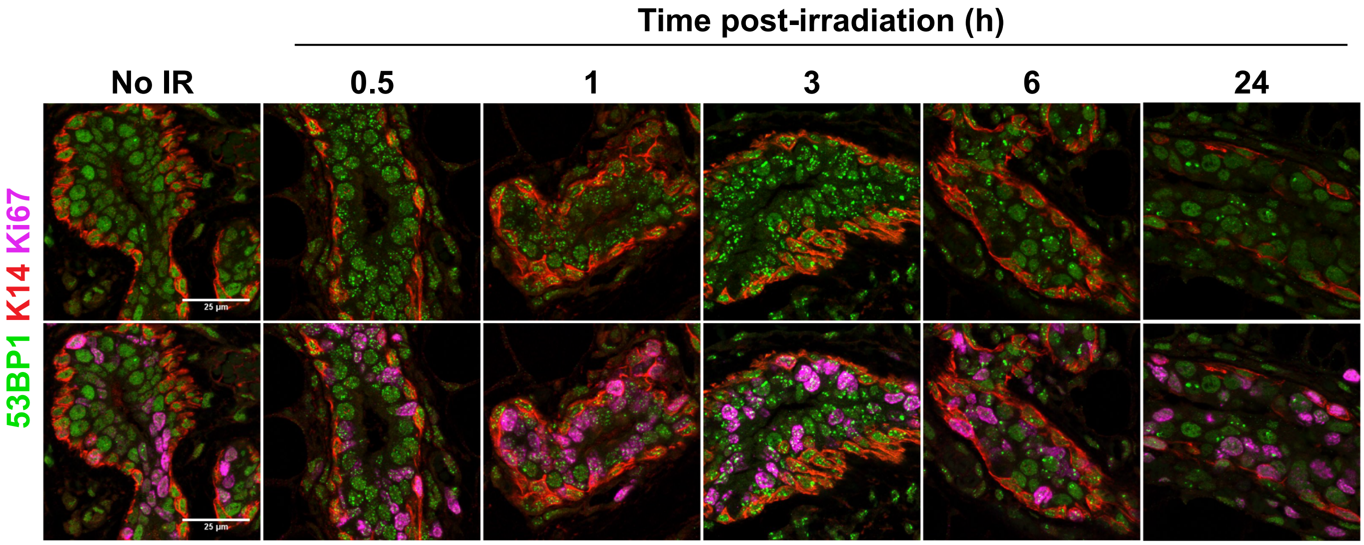

### Kim et al. Extended Data Fig. 2

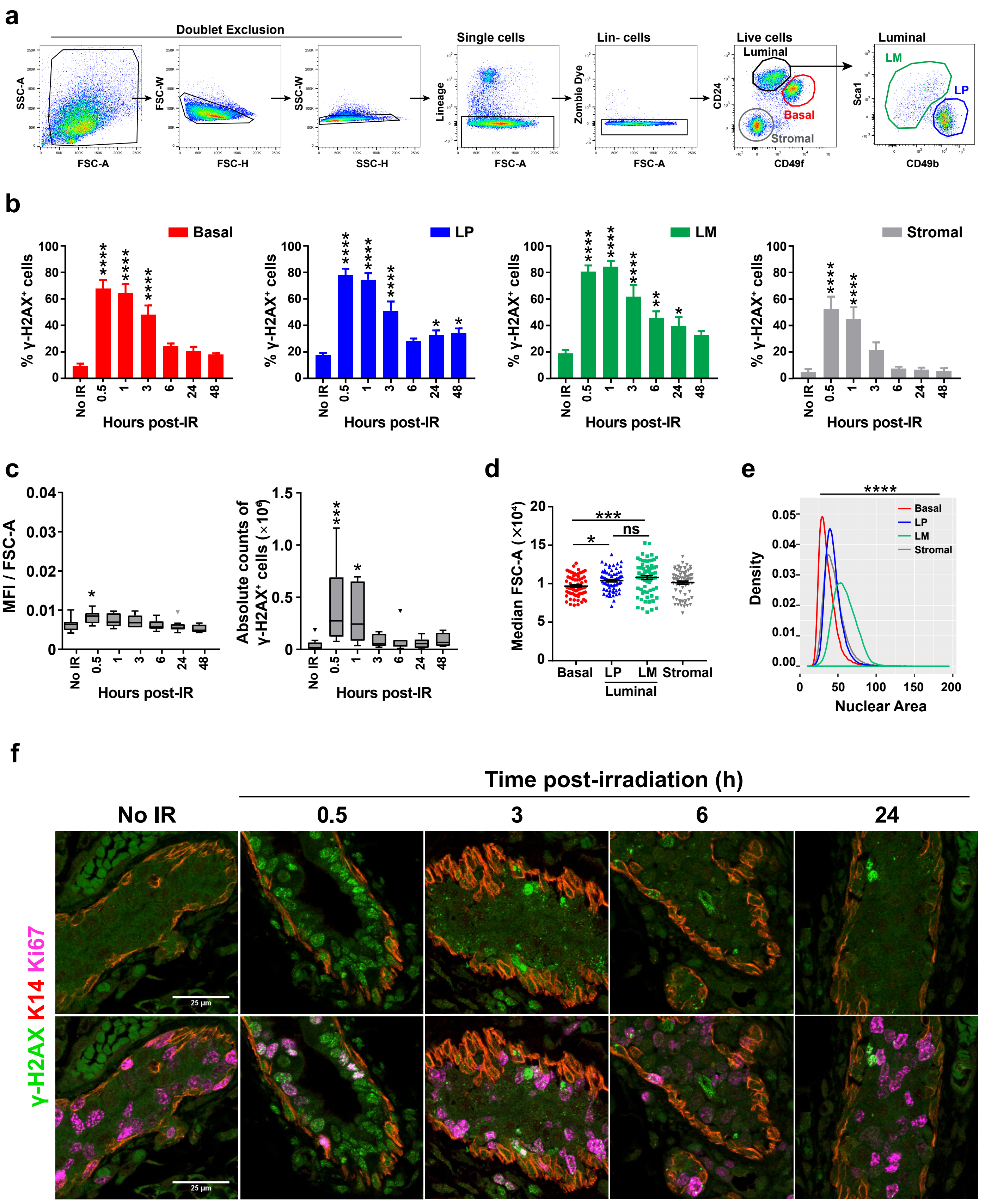

#### Kim et al. Extended Data Fig. 3

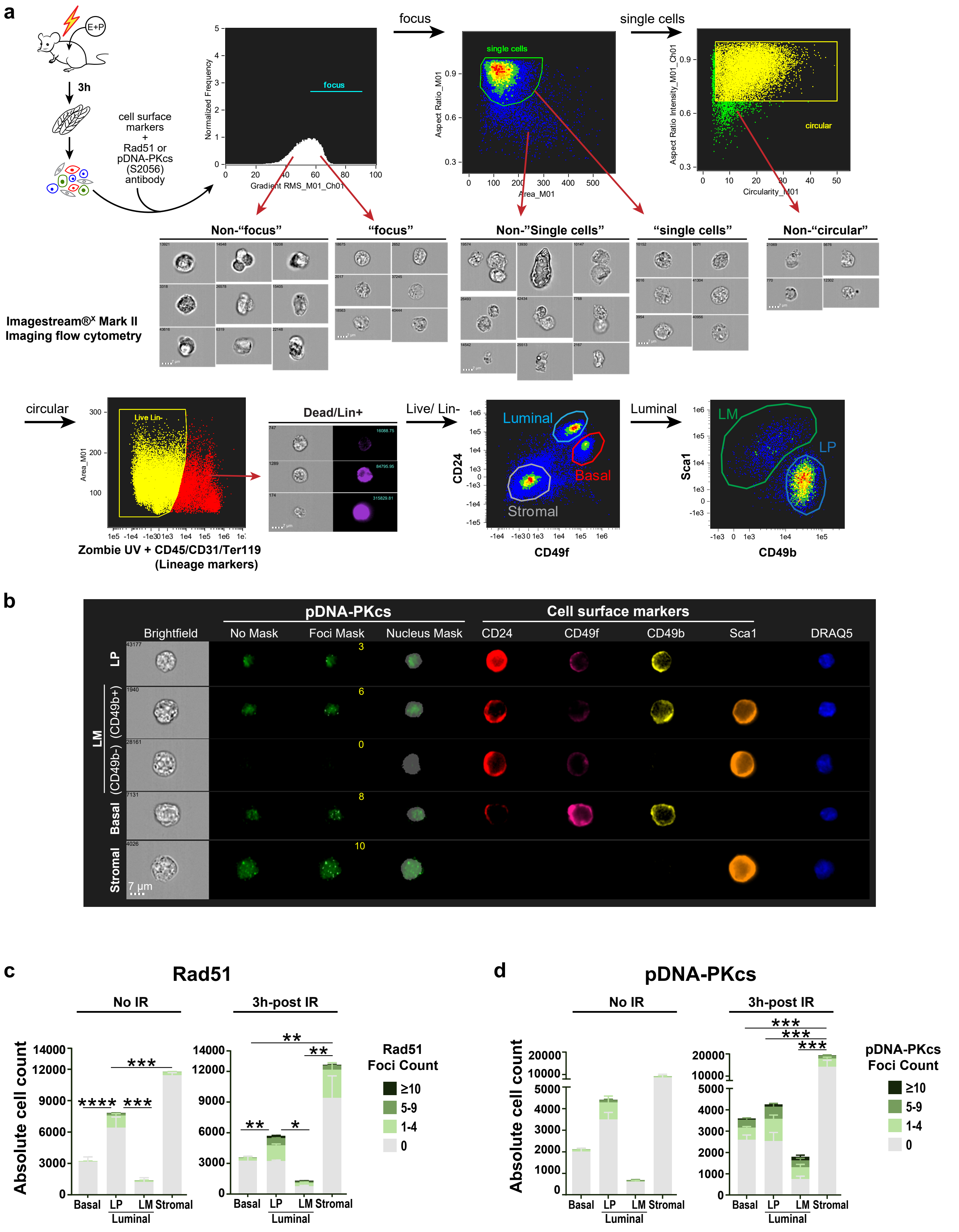

Kim et al. Extended Data Fig. 4

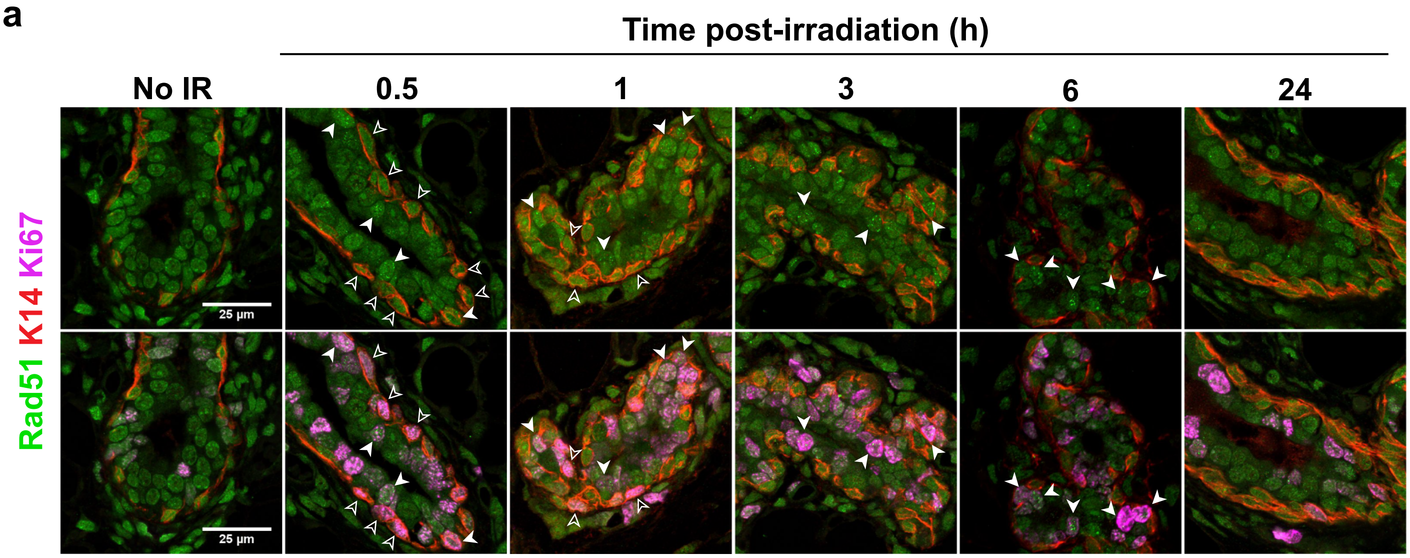


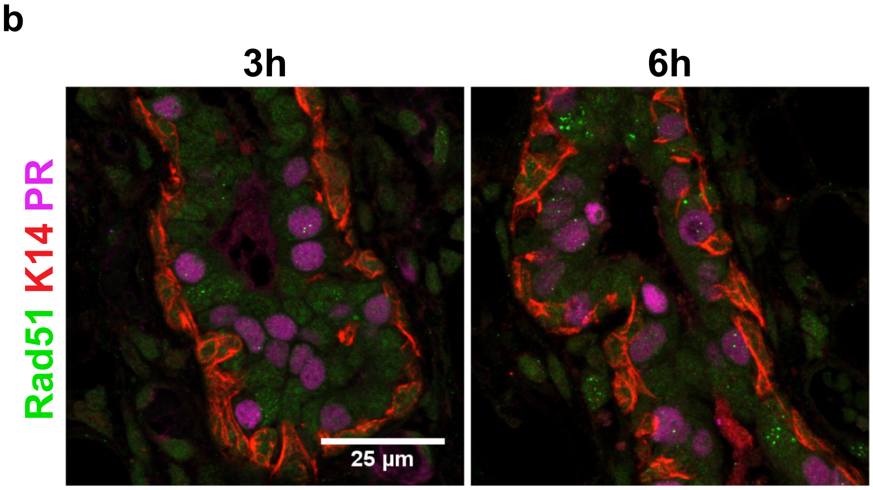


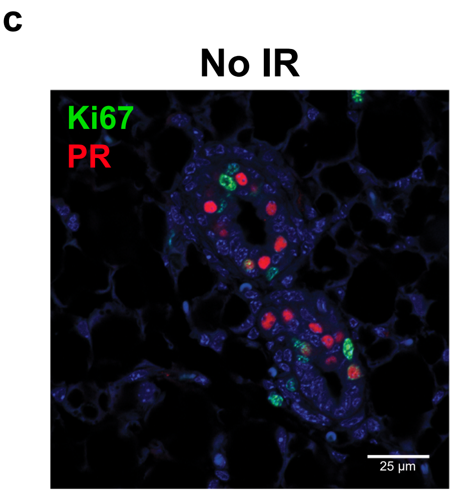

Kim et al. Extended Data Fig. 5

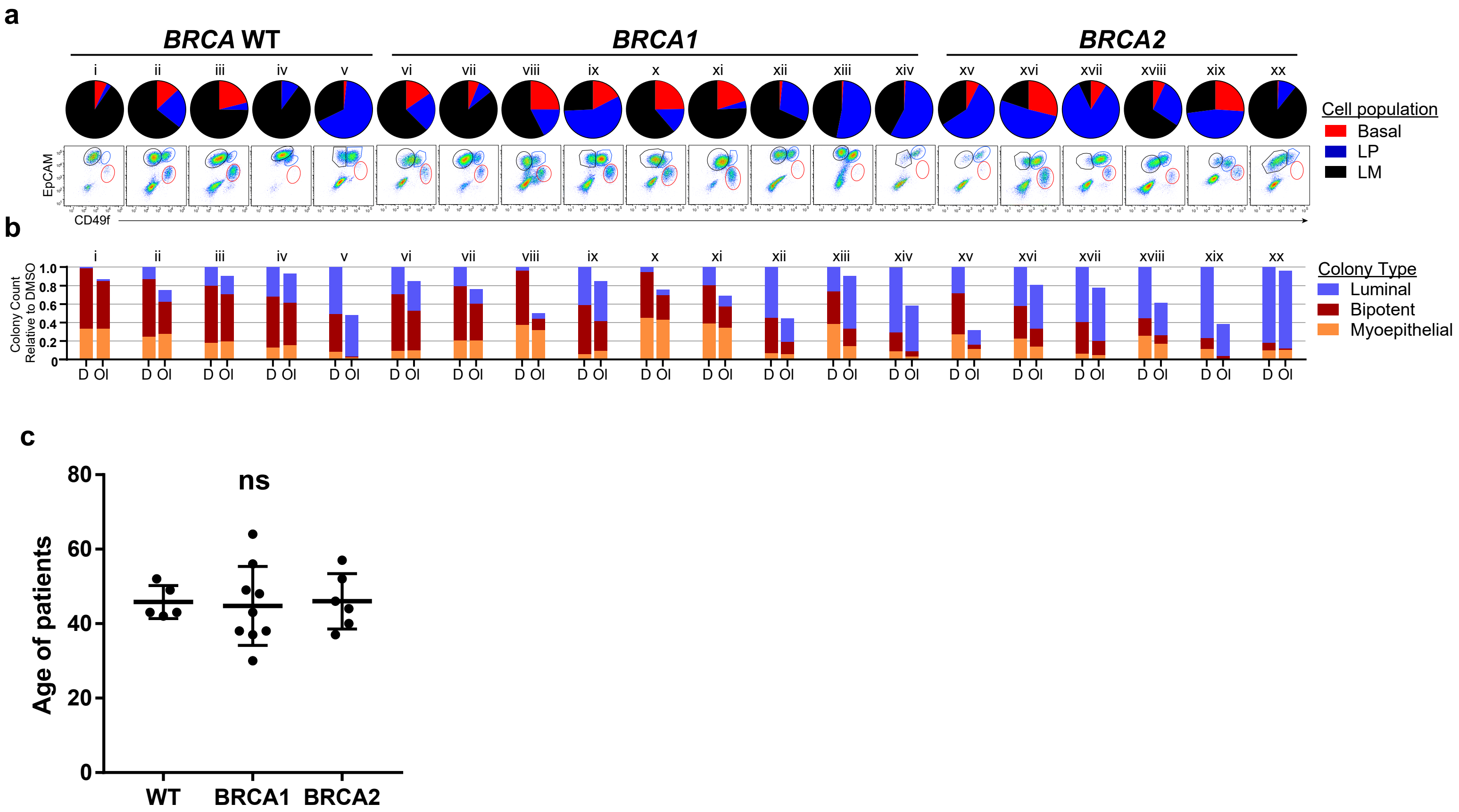

**Kim et al. Extended Data Fig. 6**

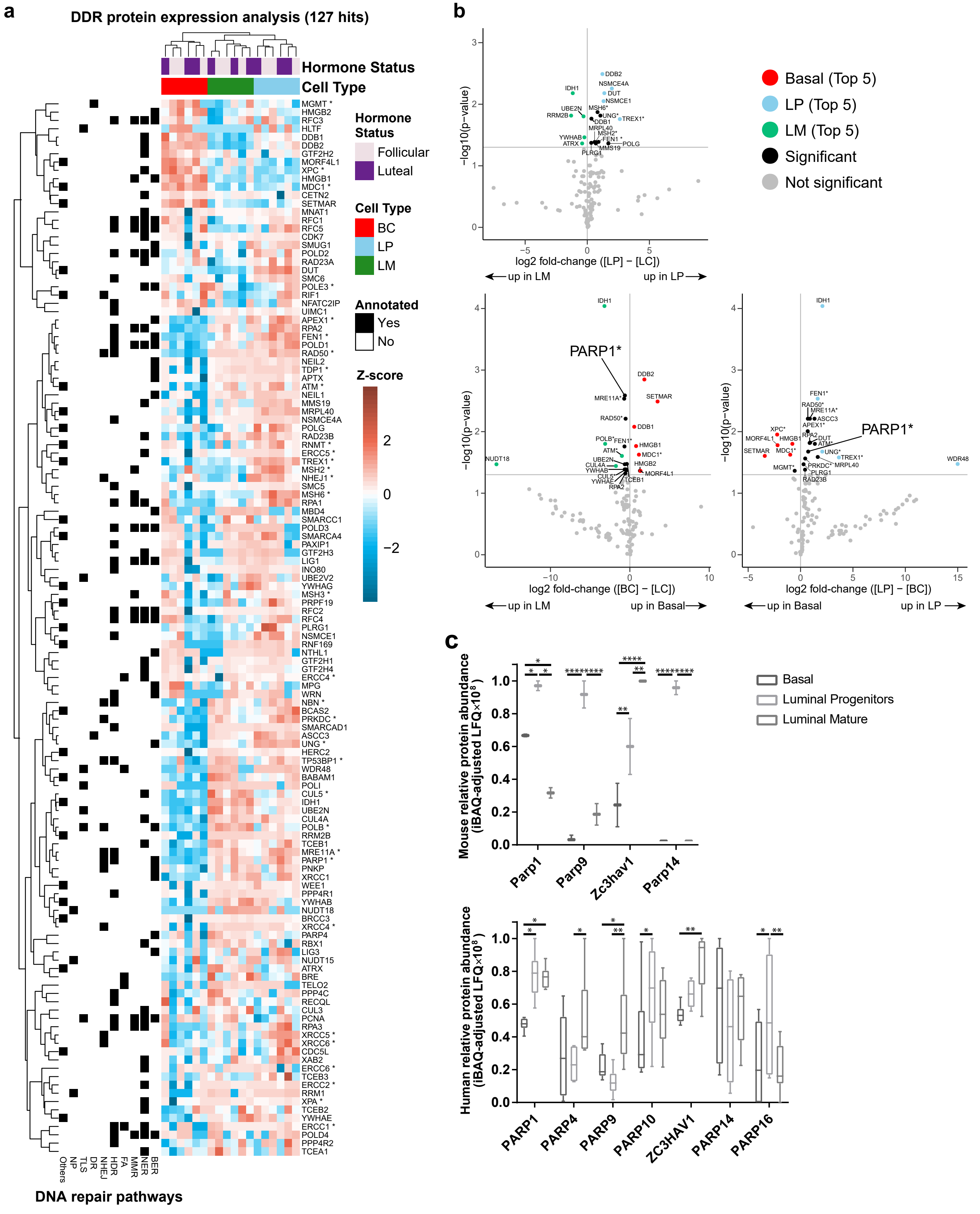

Kim et al. Extended Data Fig. 7

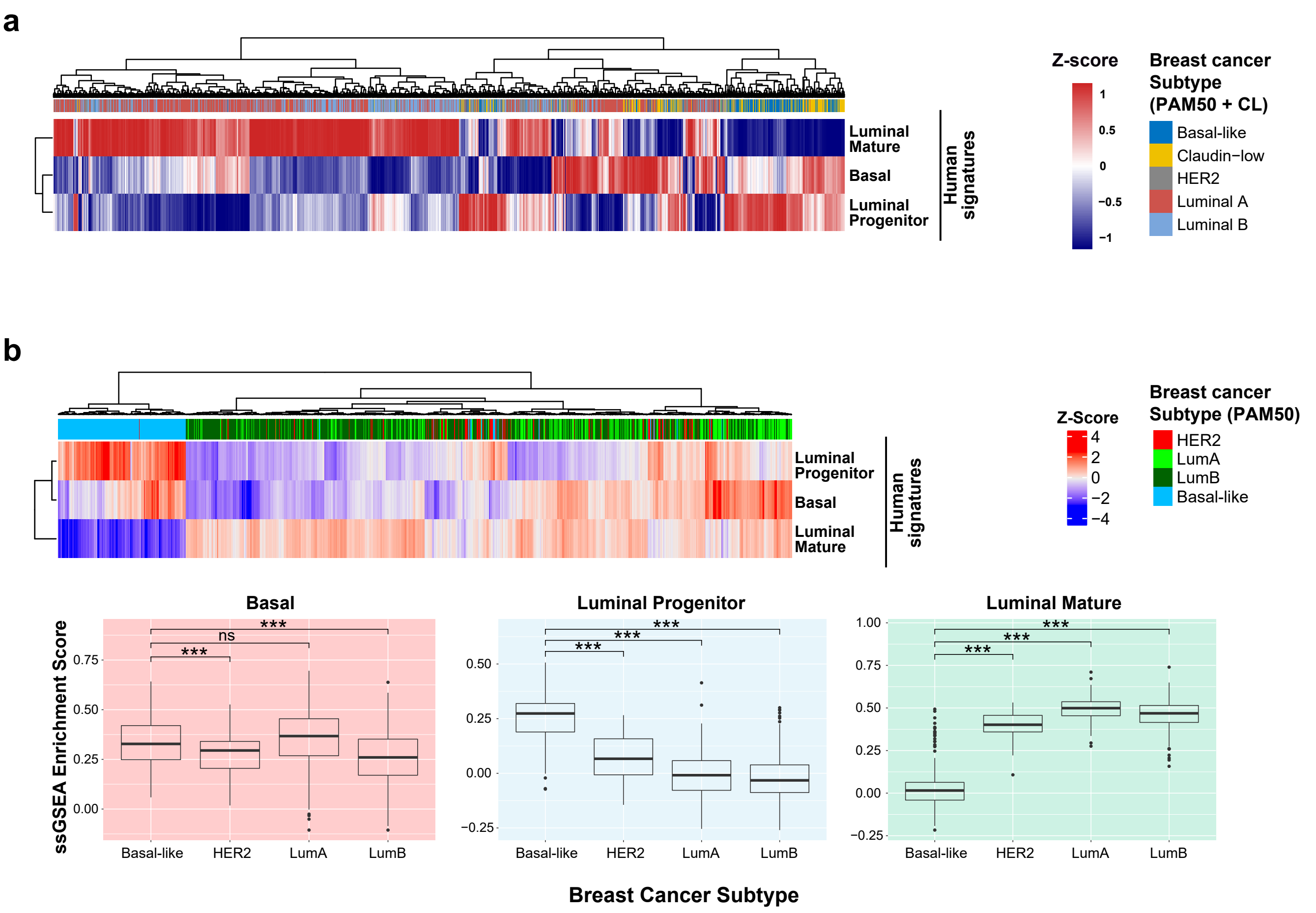

**Kim et al. Extended Data Fig. 8**

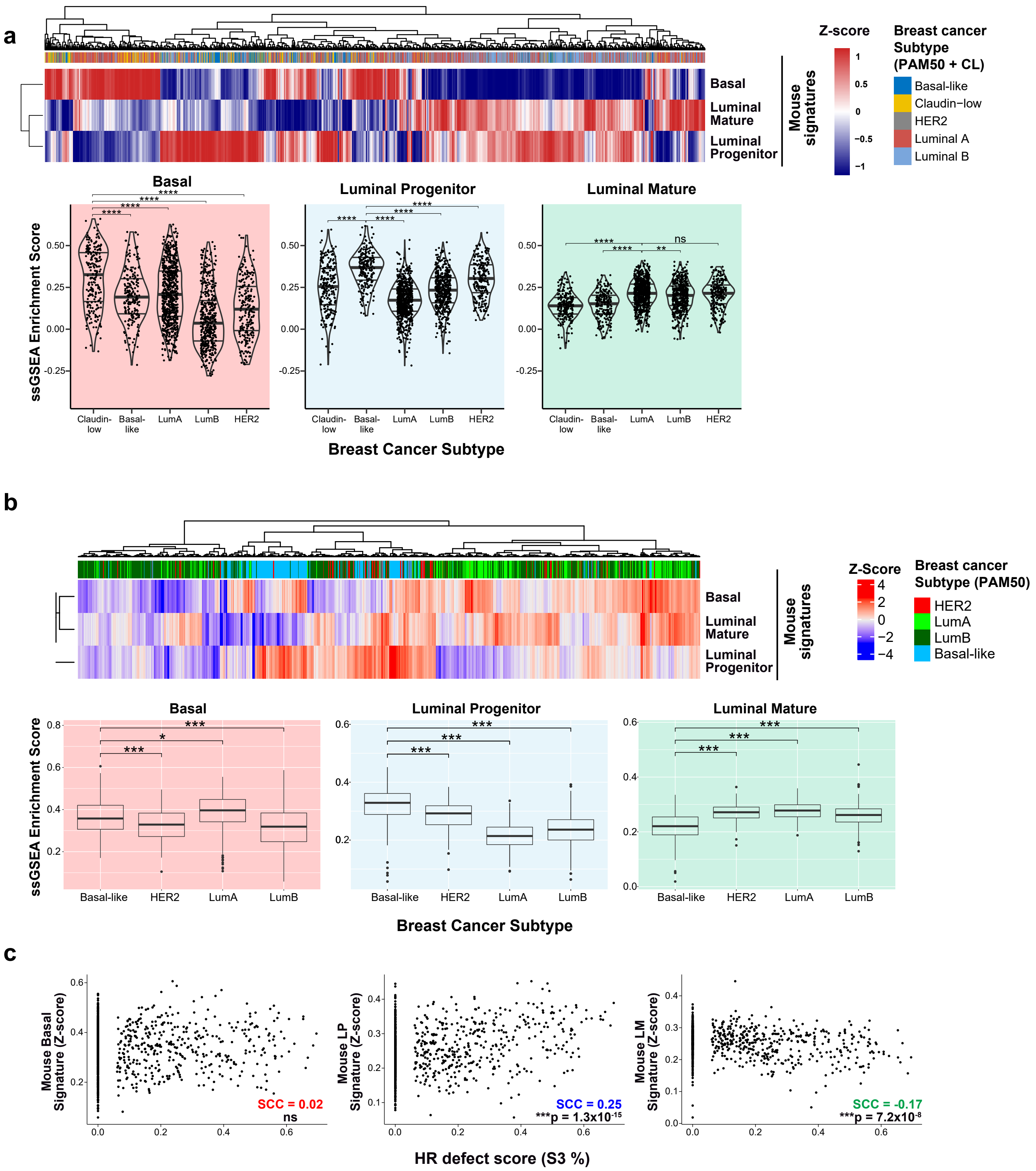

### Kim et al. Extended Data Fig. 9

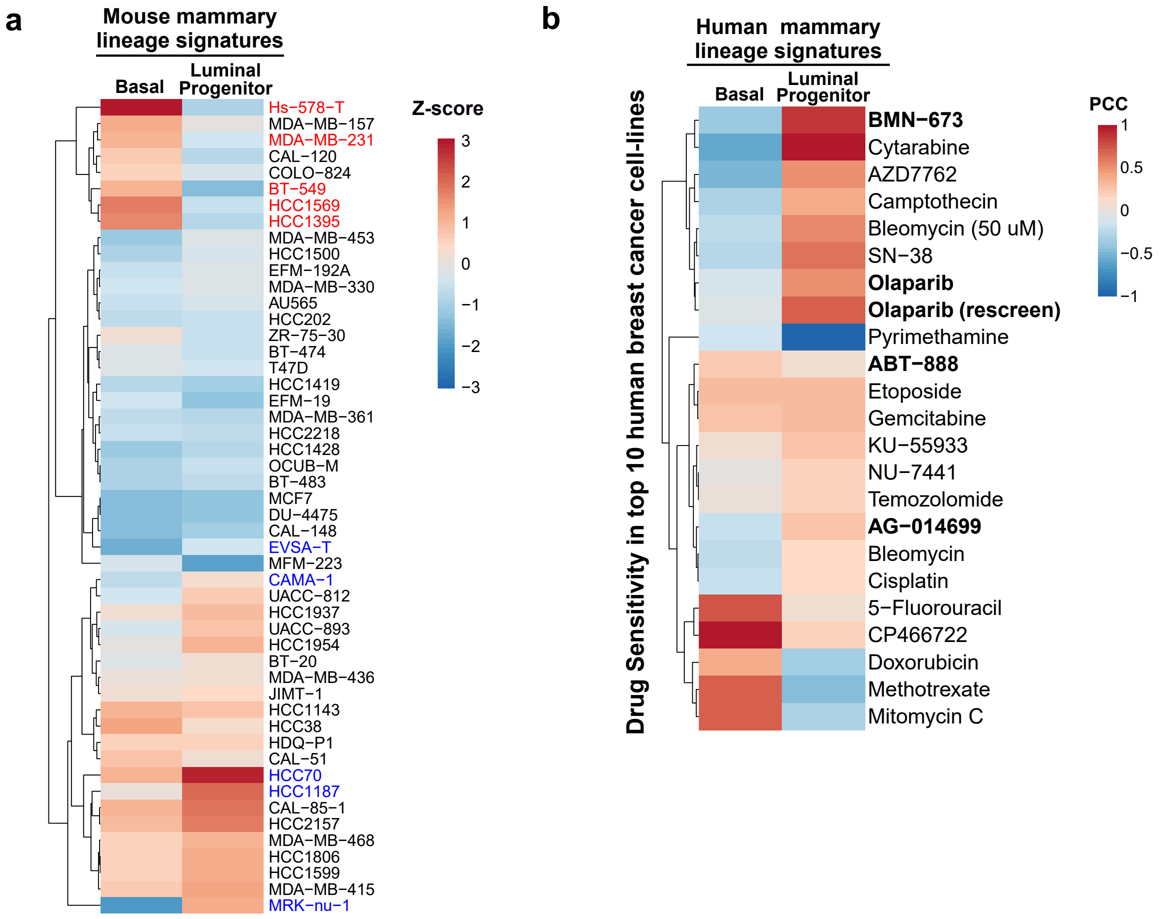

### Kim et al. Extended Data Fig. 10

**a**

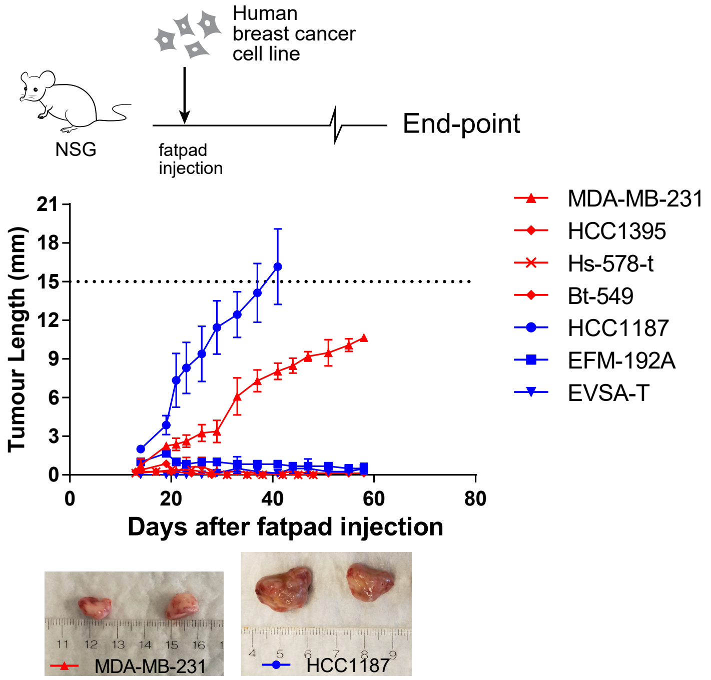


**b**

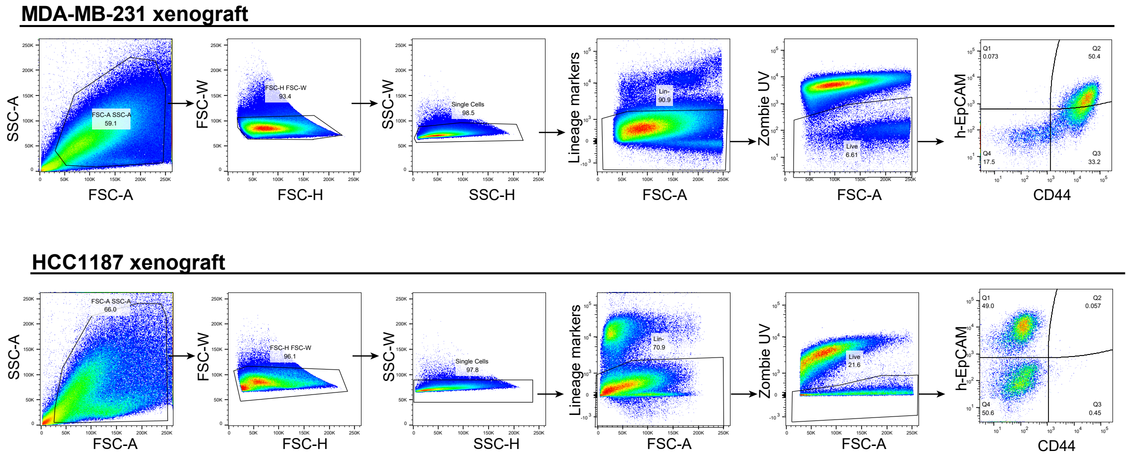
